# Multidimensional characterization of cellular ecosystems in Hodgkin lymphoma

**DOI:** 10.1101/2025.03.18.643177

**Authors:** Tomohiro Aoki, Gerben Duns, Shinya Rai, Aixiang Jiang, Andrew Lytle, Yifan Yin, Makoto Kishida, Michael Li, Denise Smorra, Laura Hilton, Stefan K Alig, Mohammad Shahrokh Esfahani, Clementine Sarkozy, Stacy Hung, Katy Milne, Adele Telenius, Luke O’Brien, Celia Strong, Talia Goodyear, Chantal Di Vito, Cassandra Luksik, Glenn Edin, Laura Gonzalez, Juan Patino Rangel, Michael Hong, Shaocheng Wu, Eric Lee, Katsuyoshi Takata, Tomoko Miyata-Takata, Merrill Boyle, Susana Ben-Neriah, Andrew P. Weng, Alexander Xu, Akil Merchant, Andrew Roth, Michael Crump, John Kuruvilla, Anca Prica, Robert Kridel, David Huntsman, Brad H Nelson, Pedro Farinha, Ryan D. Morin, Ash Alizadeh, Kerry J. Savage, David W. Scott, Christian Steidl

## Abstract

The tissue architecture of classic Hodgkin Lymphoma (CHL) is unique among cancers and characterized by rare malignant Hodgkin and Reed-Sternberg cells that co-evolve with a complex ecosystem of immune cells in the tumor microenvironment (TME). The lack of a comprehensive systems-level interrogation has hindered the description of disease heterogeneity and clinically relevant molecular subtypes. Here, we employed an integrative, multimodal approach to characterize CHL tumors using malignant cell sequencing, spatial transcriptomics and imaging mass cytometry. We identified four molecular subtypes (CST, CN913, STB, and CN2P), each characterized by distinct clinical features, mutational patterns, malignant cell gene expression profiles, and spatial architecture involving immune cell populations. Functional modeling of *CSF2RB* mutations, a characteristic feature of the CST subtype, revealed dysregulated oncogenic signaling and unique TME crosstalk. These findings highlight the significance of multi-dimensional profiling in elucidating patterns of molecular alterations that drive immune ecosystems and underlie therapeutically exploitable vulnerabilities.

## INTRODUCTION

Cancer cells exhibit heterogeneity, partially driven by molecular alterations(Andre et al., 2022; Chapuy et al., 2018; Joanito et al., 2022). With advances in DNA sequencing technology, comprehensive profiling of cancer cell genomic information has enabled the initiation of precision medicine approaches linking targeted treatments to specific molecular alterations, such as *EGFR* mutations(Maemondo et al., 2010; Paez et al., 2004). However, the systems-level description of cancer evolution in specific molecular subtypes is incomplete, as exemplified by limited knowledge about genetic and phenotypic profiles of cancer cells as part of an evolving ecosystem of malignant and non-malignant cells in the tumor microenvironment (TME). Current pathogenesis models postulate that specific malignant cell gene mutations contribute to immune escape, reflecting immune system selective pressure, and actively shape TME composition(Aoki et al., 2021b; Green et al., 2010; Steidl et al., 2011). This is especially relevant in lymphoid cancers, where various types of immunotherapies are already integral to standard treatment(Armand et al., 2018; Chen et al., 2017; Moskowitz et al., 2016; Neelapu et al., 2017; Schuster et al., 2019; Schuster et al., 2017; Timmerman et al., 2016; Younes et al., 2016).

Classic Hodgkin lymphoma (CHL) is unique among most cancers, as the malignant Hodgkin and Reed-Sternberg (HRS) cells are greatly outnumbered by reactive, non-neoplastic cells in the TME(Aoki and Steidl, 2023; Connors, 2015; Connors et al., 2020; Swerdlow et al., 2016). As such, CHL is a paradigm for a pathogenesis model that features extensive and intricate crosstalk between malignant cells and surrounding non-malignant cells(Aoki et al., 2020; Aoki et al., 2021a). The scarcity of the malignant HRS cells (estimated as 0.1-5%) has historically been a major obstacle for the study of the molecular mechanisms that underlie malignant cell crosstalk in CHL. Circulating tumor DNA (ctDNA) analysis from plasma samples partially overcomes this limitation by capturing tumor-derived somatic mutations, making it an attractive solution for clinical testing in CHL.(Alig et al., 2023; Heger et al., 2024; Reichel et al., 2015; Spina et al., 2018; Tiacci et al., 2018). Subsequently, molecular heterogeneity of CHL has been described using ctDNA technology(Alig *et al*., 2023; Heger *et al*., 2024), although functional consequences of the identified genotypes remain to be elucidated and only limited data exist on the relationship between HRS cell mutations, HRS expression phenotypes and the composition and functional state of the TME.

Based on our most recent genomic discoveries(Aoki *et al*., 2020; Aoki *et al*., 2021a; Steidl et al., 2012; Steidl *et al*., 2011; Steidl et al., 2010), we hypothesized that specific somatic gene mutations occur in response to immune pressure and alter TME architecture in CHL(Mottok and Steidl, 2015). We here sought to functionally characterize molecular heterogeneity of HRS cells directly from frozen tissue biopsies by employing HRS cell enrichment approached such as laser capture microdissection (LCM) and fluorescence-activated cell sorting (FACS)(Aoki *et al*., 2021b; Green *et al*., 2010; Maura et al., 2023; Reichel *et al*., 2015; Tiacci *et al*., 2018; Wienand et al., 2019). Furthermore, we used an integrative study design to delineate the heterogeneous molecular landscape of HRS cells and their biological contributions to form comprehensive TME ecosystems. In particular, we define a unique subset of CHL tumors with *CSF2RB* mutations, which drive deregulated, cytokine-dependent Janus tyrosine kinase (JAK)2-signal transducer and activator of transcription (STAT)5 signaling and lead to downstream immunosuppressive phenotypes. Our multimodal description of CHL heterogeneity underpins the development of a new genetic classification system, ‘HLGen’, that can serve as a biology-informed framework for testing of future precision medicine interventions.

## RESULTS

### Clinical correlates of somatic gene mutations in CHL

We analyzed tumor biopsies from 114 patients with CHL, the majority of which were obtained at time of diagnosis and treated with ABVD or ABVD equivalent therapy (n=103) **(Figure 1A)**. The median age at diagnosis was 37 years (range, 12-79) with a male predominance (58%). The baseline characteristics and outcomes in the study cohort are described in **Table S1**. We enriched HRS cells by previously established LCM(Steidl *et al*., 2010) and FACS methodology(Aoki et al., 2023; Reichel *et al*., 2015). Using these techniques, we constructed genomic libraries derived from HRS cells and applied whole exome sequencing (WES) and targeted capture sequencing **(Table S2)**. Mutational and copy number analyses identified known recurrent driver events including mutations in *SOCS1*, *STAT6*, *TNFAIP3*, *B2M* and copy number aberrations (CNAs), such as gains of 9p24.1 (including *JAK2*, *CD274* and *PDCD1LG2*, in 84% of cases), 2p15 (*REL* and *BCL11A*, 67%), and loss of 6q23.3 (*TNFAIP3,* 36%) **(Figure 1B-C, Figure S1, Table S3)**. We also identified copy number gains of 12q13.3, including focal *STAT6* amplification (30%) **(Figure 1C, Figure S2)**. These mutational findings provided a statistically powered framework to investigate the association between genomic alterations with pathological and clinical disease parameters. In particular, EBV negative (EBV-) CHL (n=77) displayed distinct genomic features compared to EBV positive (EBV+) CHL (n=25). Copy number gains of 2p15 (including *REL* and *BCL11A*) and mutations affecting *BCL7A* and *TNFAIP3* were strongly enriched in EBV- CHL **(Figure S3A and Figure S4)**. In addition, EBV- CHL cases exhibited significantly higher coding mutations overall (*p*<0.001) **(Figure S5A)**. In our cohort, the presence of anterior mediastinal disease (also referred to as “thymic” CHL)(Sarkozy et al., 2021) was observed in 47 patients (43%). Patients with thymic CHL were associated with a younger median age (27 vs. 41 years), frequent presentation with localized disease (67%), and tended to have a greater mutational burden compared to non-thymic CHL (*p*=0.077) **(Figure S5B)**. Importantly, the genomic landscape differed between thymic and non-thymic CHL cases, with *STAT6*, *GNA13*, *ITPKB*, and *TBL1XR1* mutations being significantly enriched in thymic CHL cases (*p*<0.01) **(Figure 1D)**.

**Figure 1.**
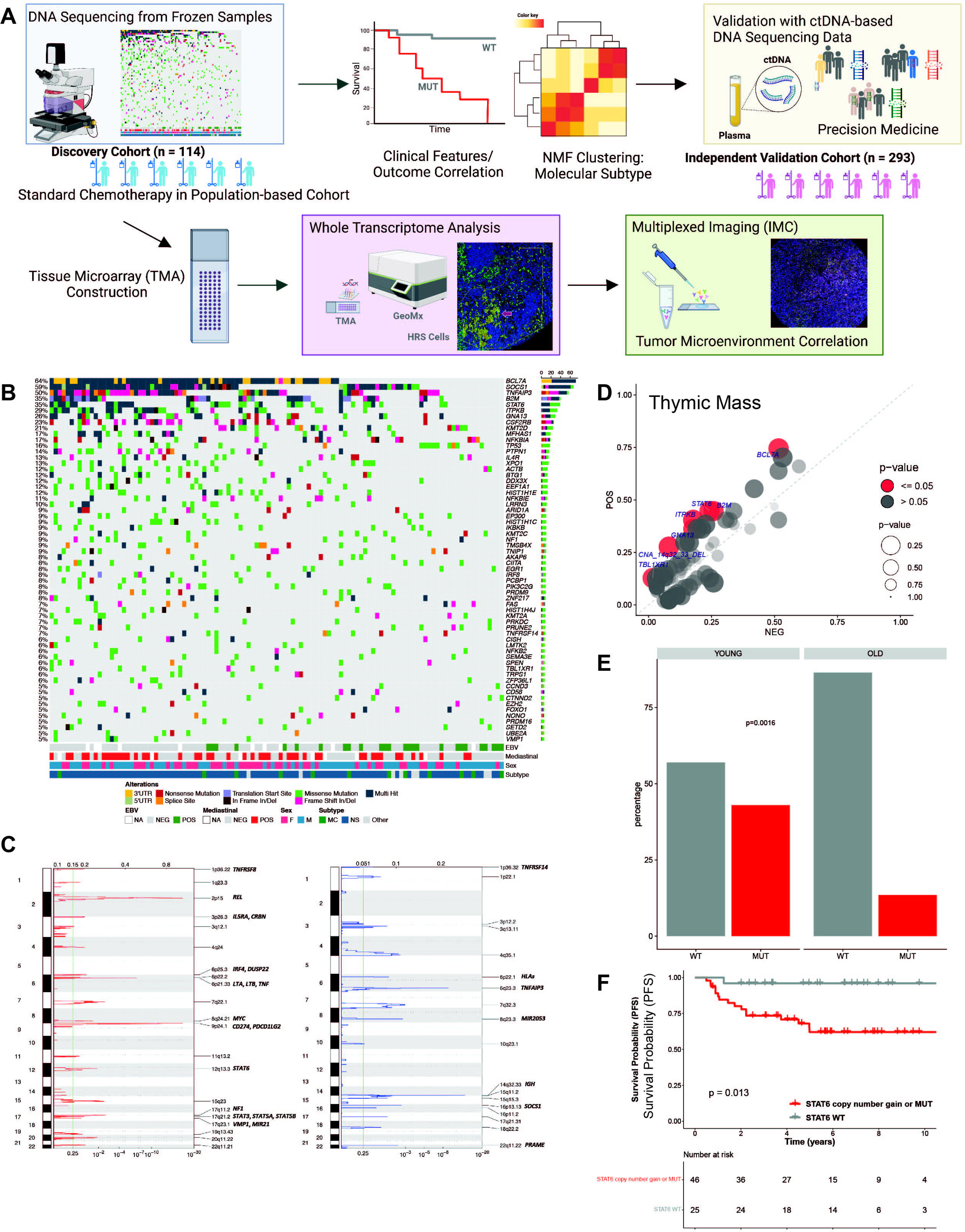
Genomic landscape and clinical correlates in CHL. (A) Cohort and study design overview. We analyzed whole exome and targeted sequencing data from 114 CHL fresh frozen samples, including 103 patients treated with ABVD-based regimens. Subsequently, we developed tissue microarrays of the cohort and applied whole transcriptome sequencing of CD30+ HRS cells using Nanostring GeoMx technology as well as multiplexed analysis using imaging mass cytometry. Key findings were validated using an independent validation cohort (n=293). (B) Oncoplot showing the most recurrently mutated genes within the CHL samples in the discovery cohort, with clinical annotations. (C) Significantly amplified (red) or deleted (blue) regions across our in-house CHL cohort. (D) Pairwise (Fisher’s exact test) comparison of the incidence of mutations within thymic vs non-thymic CHL cases. The size of the point indicating the strength of association. (E) Frequency of *STAT6* mutations in CHL according to age group (YOUNG: < 45 years, OLD: 45 years or older) at diagnosis. YOUNG: < 45 years. Fisher’s exact test was used to determine the significance of differences in observed mutational frequencies. (F) Progression-free survival (PFS) according to mutation and copy number status of *STAT6* in young CHL patients (age < 45). *P*-values were calculated using a log rank test.

Recently, Maura et al reported that pediatric and adolescents and young adult (AYA) patients (age <= 40 years) had a significantly higher mutational burden than older adults(Maura *et al*., 2023). In line with this finding, younger patients (< 45 years) in our cohort had a higher mutational burden (*p*<0.01) **(Figure S5C)**, and demonstrated a distinct genomic landscape. Mutations in *STAT6, ITPKB,* and *TNFAIP3* were significantly more frequent in younger patients(*p*<0.01), whereas mutation in *BCL2* was the most significantly enriched genetic features in older patients (*p*<0.01) **(Figure 1E and Figure S3B)**.

We next explored the association between the presence of genetic alterations and clinical outcomes. In our cohort (n=103), mutations affecting *ZNF217*, *SPEN* and *CD58* were significantly associated with poor progression-free survival (PFS) (*p*<0.001, *p*<0.01 and *p*<0.01, respectively) **(Figure S6A-D)**. Considering unique mutational patterns according to age group(Maura *et al*., 2023), we further sought to identify the genomic features specifically associated with survival outcome within these age groups. Within the group of younger patients (< 45 years), genomic alterations affecting *STAT6* (mutations or copy number gains) were most significantly associated with inferior PFS **(Figure 1F, Figure S6E, and Figure S7)**. In contrast, deletion of 4q35 was most significantly associated with poor PFS in older patients (*p*<0.001) **(Figure S8)**.

### Molecular correlates of MHC deficiency in CHL

Immune evasion by HRS cells is a hallmark of CHL and is driven by alterations affecting the antigen recognition system. Loss of major histocompatibility complex (MHC) is one of the well-known mechanisms of immunogenicity reduction(Scott and Gascoyne, 2014), and several molecular and genetic mechanisms have been reported as the basis for MHC deficiency in lymphomas(Ennishi et al., 2019; Reichel *et al*., 2015; Steidl *et al*., 2011). Here, we assessed the MHC-I and MHC-II expression status in CHL cases **(Table S3)**, as described previously(Roemer et al., 2016). Loss of MHC-I and MHC-II expression was observed in 82% and 76% of cases, respectively. MHC-I expression on HRS cells was positively correlated with EBV positivity, whereas it was negatively correlated with thymic involvement **(Figure S9A)**. Next, we investigated correlates of somatic genetic alterations with MHC status. Consistent with previous literature(Reichel *et al*., 2015), we observed an enrichment of mutations affecting *B2M*, which encodes a component of MHC-I molecules, in cases with MHC-I loss of expression (*p*<0.001) **(Figure S9B)**. In addition, mutations in *CSF2RB, GNA13*, *TNFAIP3*, and *STAT6* were significantly associated with MHC-I loss (*p*<0.001, *p*<0.001, and *p*<0.01, respectively) **(Figure S9B)**. In contrast, *CSF2RB* mutations were more frequently observed in cases with MHC-II positive HRS cells *(p*<0.01). Furthermore, we found that mutations of *PRDM9* and *LRRN*, as well as deletion of 22q11, were significantly enriched in cases with MHC-II positivity on HRS cells (*p*<0.01, *p*<0.01, and *p*<0.05, respectively) **(Figure S9C)**. Spatial transcriptomic analysis of CD30+ HRS cells using the Nanostring GeoMx platform revealed that IFNα and IFNγ signaling pathways were significantly upregulated in MHC-I positive HRS cells, consistent with transcriptional MHC-I regulation by IFNγ signaling(Zhou, 2009) **(Figure S9D-E)**. The TGFβ pathway was upregulated in MHC-II positive HRS cells **(Figure S9F)**, while a peroxisome-related gene expression signature was enriched in cases that showed loss of both MHC-I and MHC-II expression **(Figure S9G)**.

### Molecular subgroups of CHL

CHL is a clinically heterogeneous disease characterized by a bimodal age distribution, association with EBV in a subset of cases, and the presence of a bulky anterior mediastinal mass in a majority of patients. While the current pathological classification system captures some of the biological heterogeneity based on morphology and protein expression, additional features that inform on treatment strategies and prognosis are yet to be uncovered(Connors *et al*., 2020). To describe the molecular heterogeneity of HRS cells in the context of their TME ecosystem, we first applied non-negative matrix factorization (NMF) consensus clustering to a binary matrix representing the mutational or copy number status of 80 recurrently affected genes or genomic loci in our discovery cohort (n=114), and discovered four robust subgroups of tumors (clusters) with discrete genetic signatures **(Figure 2A, Figure S10)**(Brunet et al., 2004). We termed these four subgroups as CST (characteristic mutations in ***CS****F2RB* and ***T****NFAIP3*), CN913 (**c**opy **n**umber gain of **9**p24 and **13**q deletion), STB (mutations in ***ST****AT6* and ***B****2M*) and **CN**2P (**c**opy **n**umber gains in **2**p15) **(Figure 2A, Figure S10)**. Each cluster was associated with clinical features and upregulation of distinctive pathways revealed by single-sample GSEA analysis of CD30+ HRS cell whole transcriptomes (GeoMx data) **(Figure 2B-D)**. In particular, CST was characterized by younger age, and up-regulation of the STAT5 pathway **(Figure 2C-D)**. CN913 was characterized by significant upregulation of IFNγ and nuclear factor-κB (NF-κB) signaling pathways, and in 79% of cases in this cluster the HRS cells were EBV-positive (*p*<0.001, Odds Ratio 6.89) **(Figure 2B, D-E, Table S4**). STB showed significant upregulation of a TGFβ signature in HRS cells **(Figure 2D)**. CN2P included samples from mostly older patients (*p*<0.001, Odds Ratio 0.42), and showed significant upregulation of a DNA repair and a TP53 target signature **(Figure 2C-D, F, Table S4)**.

**Figure 2.**
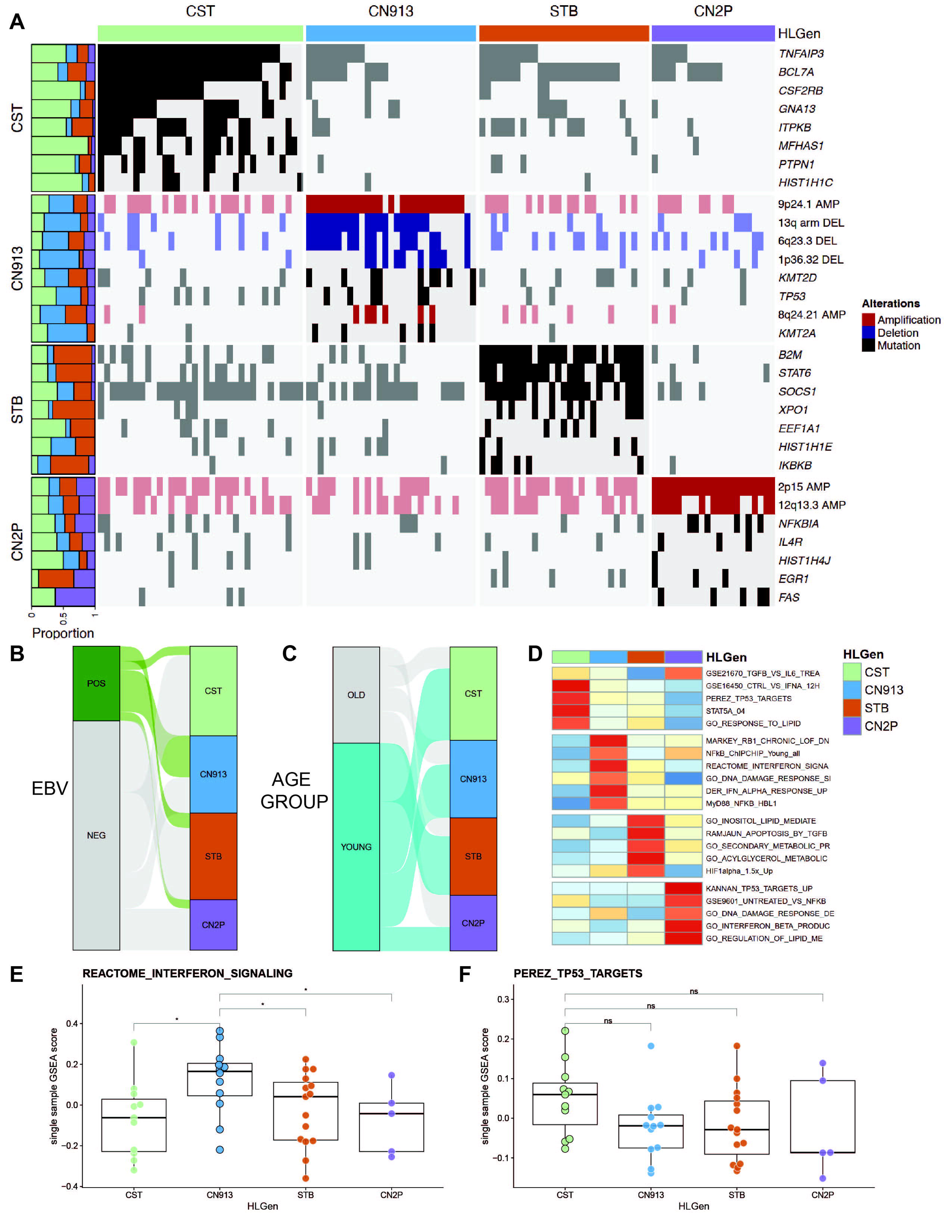
Molecular heterogeneity in CHL. (A) Non-negative matrix factorization (NMF) consensus clustering was performed using 80 molecular genomic features in our in-house cohort. Each molecular cluster is shown with their associated landmark genetic alterations. (B-C) Alluvial plots showing the correlation between molecular subtypes and EBV status of HRS cells defined by EBER IHC (B) or age (C). (D) Heatmap showing gene expression signatures enriched in CD30+ HRS cells from each molecular subtype compared to the rest of the cohort, as assessed by whole transcriptome analysis using the Nanostring GeoMx platform. (E-F) ssGSEA scores for interferon signaling (E) and a TP53 target signature (F) across HLGen subtypes. Statistical analyses were conducted using one-way ANOVA followed by Dunnett’s multiple comparisons test.

### Molecular subgroups are linked to the spatial architecture of the TME

We next sought to understand the impact of molecular alterations on the TME ecosystem. To delineate the spatially resolved TME architecture according to HRS cell mutational profiles, we applied imaging mass cytometry (IMC) to the intact tissue biopsies of the exome/targeted sequencing cohort, arranged on tissue microarrays. The IMC panel was designed to inform on key cellular interactions discovered in previous studies using single cell (sc) RNA-sequencing (RNA-seq) **(Figure S11, and Table S5)**(Aoki *et al*., 2020; Aoki *et al*., 2021a). To capture spatial patterns in the TME, we calculated the “spatial score”, an algorithm previously established to measure the distance of nearest neighbor cells to the malignant CD30+ HRS cells within close interaction range(Aoki et al., 2024) **(Figure S11)**. IMC analysis identified correlations between each molecular subgroup and TME composition, in particular with specific CD4+ T cell subsets. The CST subgroup showed a significant enrichment of FOXP3+ regulatory T cells (Tregs) among other molecular subgroups, the CN913 subgroup a significant enrichment with LAG3+CD4+ T cells and the CN2P subgroup a significant enrichment of PD1+CD4+ T cells **(Figure 3A-B and Figure S12A)**. The observed correlations between molecular subtypes and TME composition are consistent with, and significantly expand upon, a previously reported negative correlation between FOXP3+ Tregs and LAG3+ Tregs by our group(Aoki *et al*., 2020). We further identified a significant spatial enrichment of a PDL1+ macrophage population surrounding HRS cells in CN913, where PDL1 copy number gains in HRS cells were frequent **(Figure 3B)**. In contrast, in the STB subgroup, which is characterized by the presence of *B2M* mutations, a GZB+ cytotoxic CD8+ T cell population was less frequent indicating an immune-cold microenvironment **(Figure 3B)**. We next applied the cell-to-cell communication inference tool, Cell Chat(Jin et al., 2021), and confirmed previously reported interactions between CXCR5+ HRS cells and CXCL13+ macrophages(Aoki *et al*., 2024) in CN913 **(Figure S12B)**, and between PD-L1+ HRS cells and PD1+CD4+ T cells(Yamamoto et al., 2008) in CN2P **(Figure 3C)**. Additionally, we identified novel associations including interaction of CCL17 (also known as Thymus and Activation-Regulated Chemokine, TARC)- positive HRS cells with CCR4+CD4+ T cells in CST and Galectin-9+ HRS cells with TIM3+ macrophages in STB **(Figure 3C and Figure S12B)**. We validated the TARC+ HRS cell and CCR4+CD4+ T cell interaction using the iTALK method(Wang et al., 2019) **(Figure 3D)** and directly visualized the presence of CCR4+CD4+ T cells in close proximity to TARC+ HRS cells in the IMC data (**Figure 3E-F).** Importantly, each significant cell-to-cell interaction was only observed within each distinct molecular subgroup **(Figure 3C)**.

**Figure 3.**
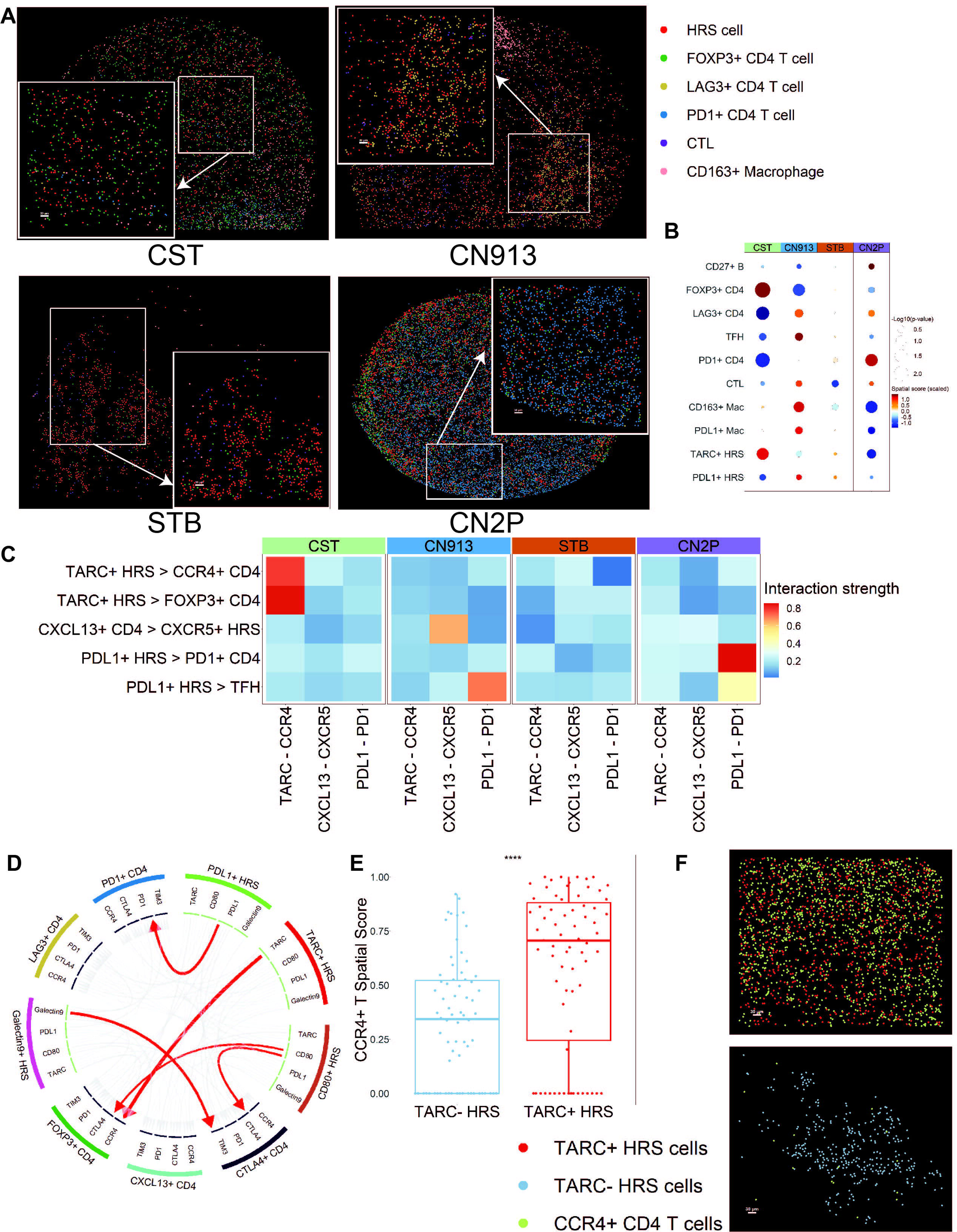
Tumor-microenvironment ecosystems linked to molecular subtypes in CHL. (A) Membrane map depicting major immune cell populations in each molecular subgroup. The CST subgroup (top left) showed a significant enrichment of FOXP3+ regulatory T cells (Tregs) (green) among other molecular subgroups, the CN913 subgroup (top right) a significant enrichment with LAG3+CD4+ T cells (yellow) and the CN2P subgroup (bottom right) a significant enrichment of PD1+CD4+ T cells. (blue), In contrast, cytotoxic T cell (CTL) population (purple) was less frequent in STB (bottom left), indicating an immune-cold microenvironment. (B) Spatial score for the indicated immune cell population by imaging mass cytometry (IMC) according to molecular subgroup. Dot plot showing correlation of spatial scores of major immune cell and HRS cell markers with molecular subtypes by IMC. (C) Heatmap shows significant ligand and receptor interactions between HRS cells and CD4+ T cell populations using CellChat in each molecular subtype. (D) An interaction between CD4+ T cells and HRS cells in CHL samples was predicted using the iTALK tool. (E) Boxplot showing the spatial score of CCR4+ T cells in the region surrounding TARC+ HRS cells or TARC- HRS cells. An unpaired, two-sided *t*-test was used to compare the two conditions. (F) Membrane map depicting CCR4+CD4+ T cells (light green), TARC+ HRS cells (red) and TARC- HRS cells (light blue).

### C-terminal truncating *CSF2RB* mutations promote cytokine-dependent gain-of-function phenotypes

CSF2RB, also known as the β common chain receptor (βc), is a component of cytokine receptors with specificity to IL-3, IL-5, and GM-CSF, which activate signaling pathways, such as JAK2-STAT5(Dougan et al., 2019). Considering the high mutation frequency of *CSF2RB* in CST and their high specificity to CHL in contrast to other major lymphoma entities(Pastore et al., 2015; Reddy et al., 2017; Schmitz et al., 2018; Wienand *et al*., 2019), we hypothesized that *CSF2RB* mutations contribute to dysregulated oncogenic signaling, leading to both cell-autonomous and TME-related phenotypes in CHL. Notably, *CSF2RB* exhibited the highest proportion of truncating mutations among all evaluated genes **(Figure 4A)**. In our CHL cohort, we identified *CSF2RB* mutations in 26 of 107 cases (24.3%), mostly affecting the cytoplasmic domain (exon 14). These mutations include missense (n=2), multiple-hit (n=3), and truncating mutations (n=21), with the most frequent being E788* hotspot mutations **(Figure 4B)**.

**Figure 4.**
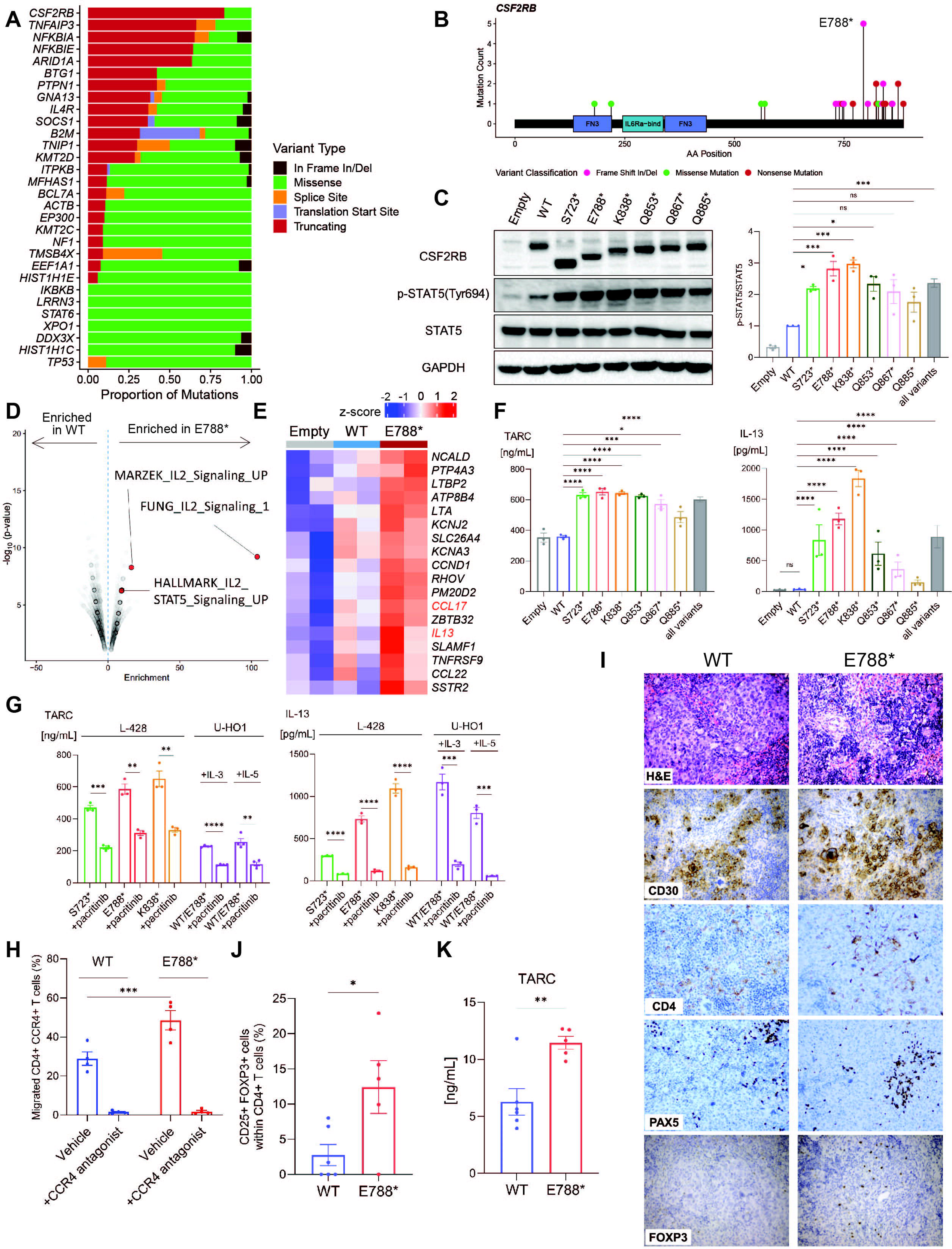
Functional characterization of *CSF2RB* mutations *in vitro* and *in vivo*. (A) Proportion of mutation types across recurrently mutated genes. (B) Lollipop diagram depicting the distribution of *CSF2RB* mutations in our in-house cohort. (C) WB analysis of CSF2RB expression and phosphorylation status of STAT5 in the L-428 model. Representative images are shown (left). Densitometry analysis was carried out by Image Lab (right). Data are presented as mean ± SEM. Statistical analyses were conducted using one-way ANOVA followed by Dunnett’s multiple comparisons test. (D) Top pathways from LymphoChip upregulated in E788* cells compared to WT cells in the L-428 model. (E) Heatmap showing transcript levels of genes in the L-428 model. Genes with consistent expression changes (absolute fold change ≥ 1.4) across all three experimental models (L-428, U-HO1 + IL-3, U-HO1 + IL-5) are shown. (F) TARC and IL-13 concentrations in supernatant of L-428 model at baseline. Data are presented as mean ± SEM. Statistical analyses were conducted using one-way ANOVA followed by Dunnett’s multiple comparisons test (TARC), and a negative binomial regression model to account for overdispersion in the data (IL-13). (G) TARC and IL-13 concentrations in supernatant of L-428 (left) model treated with pacritinib (0.5 μM) for 48 hours, and U-HO1 (right) model in the presence of IL-3 (1 ng/mL) or IL-5 (1 ng/mL) treated with or without pacritinib (0.5 μM) for 48 hours. Data are presented as mean ± SEM. Pre- and post-treatment comparisons were conducted using an unpaired, two-sided *t*-test for each cell type. (H) CCR4+CD4+ T cell migration towards L-428 CSF2RB WT or E788* supernatant in a transwell migration assay, showing the percentage of migrated cells relative to the input control. A paired, two-sided *t*-test was used to compare the two conditions. Data are presented as mean ± SEM (n=4). (I) Spleen tissues from humanized mouse model, sacrificed at 12 weeks after CB injection, were stained with hematoxylin and eosin (H&E), CD30, CD4, PAX5, and FOXP3. The image is presented at a magnification of 20X. (J) Percentage of CD4+CD25+FOXP3+ Tregs identified in spleen of humanized CNW mice engrafted with L-428 CSF2RB WT or E788* cells by FCM. An unpaired, two-sided *t*-test was used to compare the two conditions, and data are shown as the mean ± SEM (n=6 for WT and n=5 for E788*). (K) Bar graph depicts serum TARC concentration from CNW mice engrafted L-428 WT (n=6) and E788* (n=5) cells, sacrificed at 12 weeks after CB injection. An unpaired, two-sided *t*-test was used to compare the two conditions. The data are shown as mean ± SEM.

To functionally characterize the *CSF2RB* mutations observed in CHL, we selected the Hodgkin lymphoma-derived cell line, L-428, to generate isogenic CSF2RB overexpression systems. Specifically, to assess the full spectrum of mutations affecting the cytoplasmic domain, we stably transduced L-428 cells with mutant *CSF2RB* constructs, including S723*, E788*, K838*, Q853*, Q867*, and Q885*, ensuring equivalent transcript levels as confirmed by qPCR **(Figure S13A)**. Western blot (WB) analysis confirmed that all mutations resulted in premature CSF2RB protein truncation **(Figure 4C)**. Next, we assessed the impact of the introduced CSF2RB truncating mutations on downstream signaling in the L-428 model, in which IL-5 is produced in an autocrine manner **(Figure S13B)**. WB and FCM analyses showed that STAT5 was hyperphosphorylated in all truncating mutants compared to non-transduced (Empty) and WT cells, with no significant changes observed in the activation of other STAT proteins **(Figure 4C and Figure S13C-D)**. Importantly, STAT5 hyperactivation was inhibited by an antibody targeting surface IL5RA **(Figure S13E)**, suggesting that the C-terminal truncating *CSF2RB* mutations promote cytokine-dependent gain-of-function phenotypes.

For validation of the findings in L-428 cells, we also generated a U-HO1 model with *CSF2RB* heterozygous genotypes by 1:1 transduction of WT and mutant alleles (WT/ E788*) **(Figure S14A)**. Upon IL-3 or IL-5 stimulation, we observed markedly higher p-STAT5 induction with E788* and WT/E788* compared to WT in U-HO1 cells, while no differences were detected at baseline or after GM-CSF stimulation, validating the cytokine-dependent gain-of-function phenotypes **(Figure S14B)**. To assess the effect of the *CSF2RB*-E788* mutation on the HRS transcriptome, we applied RNA-seq to both L-428 (Empty vs WT vs E788*) and U-HO1 (WT vs WT/ E788*) models. Differential gene expression analysis (DGEA) followed by gene ontology (GO) analysis revealed upregulation of IL-2 signaling pathway-related genes in E788* cells compared to WT cells in the L-428 model **(Figure 4D and Table S6)**, which was validated in the U-HO1 models under IL-3 or IL-5 stimulation **(Figure S14C and Table S7-8)**. Additionally, we identified 19 genes with consistent expression changes across all three experimental models (L- 428, U-HO1+IL-3, U-HO1+IL-5) (absolute fold change ≥ 1.4) **(Figure S14D and Table S9)**, including *CCL17* and *IL-13* **(Figure 4E)**, both of which are key cytokines/chemokines produced by HRS cells(Alig *et al*., 2023). In accordance with the DGEA results, we detected higher amounts of TARC and IL-13 by Enzyme-Linked Immunosorbent Assay (ELISA) in the supernatant from cells expressing all truncating mutants compared with WT cells in the L-428 model, and both experimental U-HO1 models (WT vs WT/E788*) **(Figure 4F and Figure S14E)**. Furthermore, these gain-of-function phenotypes were reversed by a selective JAK2 inhibitor (pacritinib)(Derenzini and Younes, 2013) **(Figure 4G and Figure S14F-G)**, confirming the specificity of increased TARC and IL-13 protein expression by cytokine-dependent, deregulated signaling through the JAK2-STAT5 axis.

### *CSF2RB* mutation-associated changes in TME composition and cellular crosstalk

TARC interacts with CCR4, a receptor highly expressed on Tregs, mediating CCR4+ cell chemotaxis(Sugiyama et al., 2013). To evaluate if the increased TARC production in E788* cells enhances CCR4+ T cell migration, we performed transwell assays with CCR4+ cell populations, including Tregs, isolated from peripheral blood mononuclear cells (PBMCs) of four different healthy donors **(Figure S15A-B)**. Supernatant from E788* cells significantly promoted CCR4+ T cell migration compared to WT (*p*<0.001), and this effect was abrogated by a CCR4 antagonist **(Figure 4H)**. Next, we investigated whether the increased TARC production influences the TME in a humanized xenotransplantation model, co-engrafted with human CD34+ cord blood (CB) cells and either L-428 CSF2RB-WT (WT group) or E788* (E788* group) cells **(Figure S15C)**. We analyzed mice with comparable CD45+ cell engraftment and similar tumor burden, measured by in vivo imaging system, to ensure consistency between WT and E788* groups **(Figure S15D-E)**. Histological analysis revealed scattered large CD30+ L-428 cells in both groups, surrounded by infiltrating human CB-derived CD4+ T cells, reminiscent of the patterns commonly observed in the TME of primary CHL samples **(Figure 4I)**. Notably, and in line with the TME features of the CST subtype, CD4+CD25+FOXP3+ Tregs infiltration was significantly increased in the E788* group compared to the WT group (*p*<0.05), while no major differences were observed in the overall proportions of CD19+ B-cells, CD4+ T cells and CD8+ T cells **(Figure 4J and Figure S15F)**. We also confirmed significantly higher serum TARC levels in the E788* group compared to the WT group (*p*<0.01) **(Figure 4K)**. Taken together, we demonstrate that the gain-of function phenotype of *CSF2RB* truncating mutations contributes to an increased Treg infiltration in the TME through the TARC-CCR4 axis in CHL.

### Development of a CHL molecular classification system

Using the CSF2RB *in vitro* and *in vivo* models and clinical associations, we confirmed that molecular alterations drive the generation of distinct HRS cell-autonomous and TME-related phenotypes in CHL, contributing to biological heterogeneity. The strong association between genetic subgroups and distinct TME ecosystems suggests opportunities for immunotherapeutic targeting. Therefore, we aimed to establish a molecular classification system that could reliably assign patient tumors to our NMF-based molecular subgroups using mutational profiles **(Figure 5A)**. To achieve this goal, we developed a molecular classification tool, “HLGen”, leveraging and integrating machine-learning outputs from 11 distinct classification algorithms. HLGen was trained on our NMF-based molecular clusters using the 80 genomic features that were originally used for cluster assignment **(Training cohort, Figure 2A**, **Figure 5A, Figure S16, Table S2 and STAR ⍰ METHODS)**. Inclusion of all features at the stage of classifier development makes the classifier applicable to a broader set of datasets generated with various targeted sequencing panels. Algorithms were selected for their complementary characteristics, with each algorithm independently trained on our training cohort to ensure high accuracy between NMF clusters and HLGen classification (> 85%) and generalizability **(Table S10)**. To verify the stability of our molecular classification system and to validate cluster correlates with clinical features, we applied HLGen to somatic variant and copy number calls from an independent CHL cohort of 293 patients that were obtained by targeted sequencing of ctDNA(Alig *et al*., 2023) **(“validation cohort”, Figure S17)**. We reproduced the four molecular classes irrespective of the difference in the sequencing pipeline with common mutations and copy number aberrations which contribute to defining each cluster **(Figure 5B-C and Table S11-12)**. Furthermore, in line with our training cohort, the clinically significant correlations of the CN913 cluster with EBV positivity on HRS cells (*p*<0.001) and CN2P with age (*p*<0.0001), were observed in the validation cohort **(Figure 5D-E and Table S4)**, confirming the biological robustness of HLGen. Additionally, HLGen recognizes a group of unclassified cases, for which confident assignment cannot be made to any of the molecular subtypes. Among the 293 cases in the validation cohort, 51 (17%) cases were assigned as unclassified **(Table S12)**. When comparing the molecular assignment of HLGen with the H1/H2 classification proposed by Alig et al(Alig *et al*., 2023), both methods identified classes with lower tumor mutational burden (CN913 and CN2P in HLGen, and H2 in the validation study), which were largely correlated to each other **(Figure 5F)**. HLGen is available as an online tool (https://shiny.bcgsc.ca/HLGen/) for further validation in routine clinical practice and future clinical trials.

**Figure 5.**
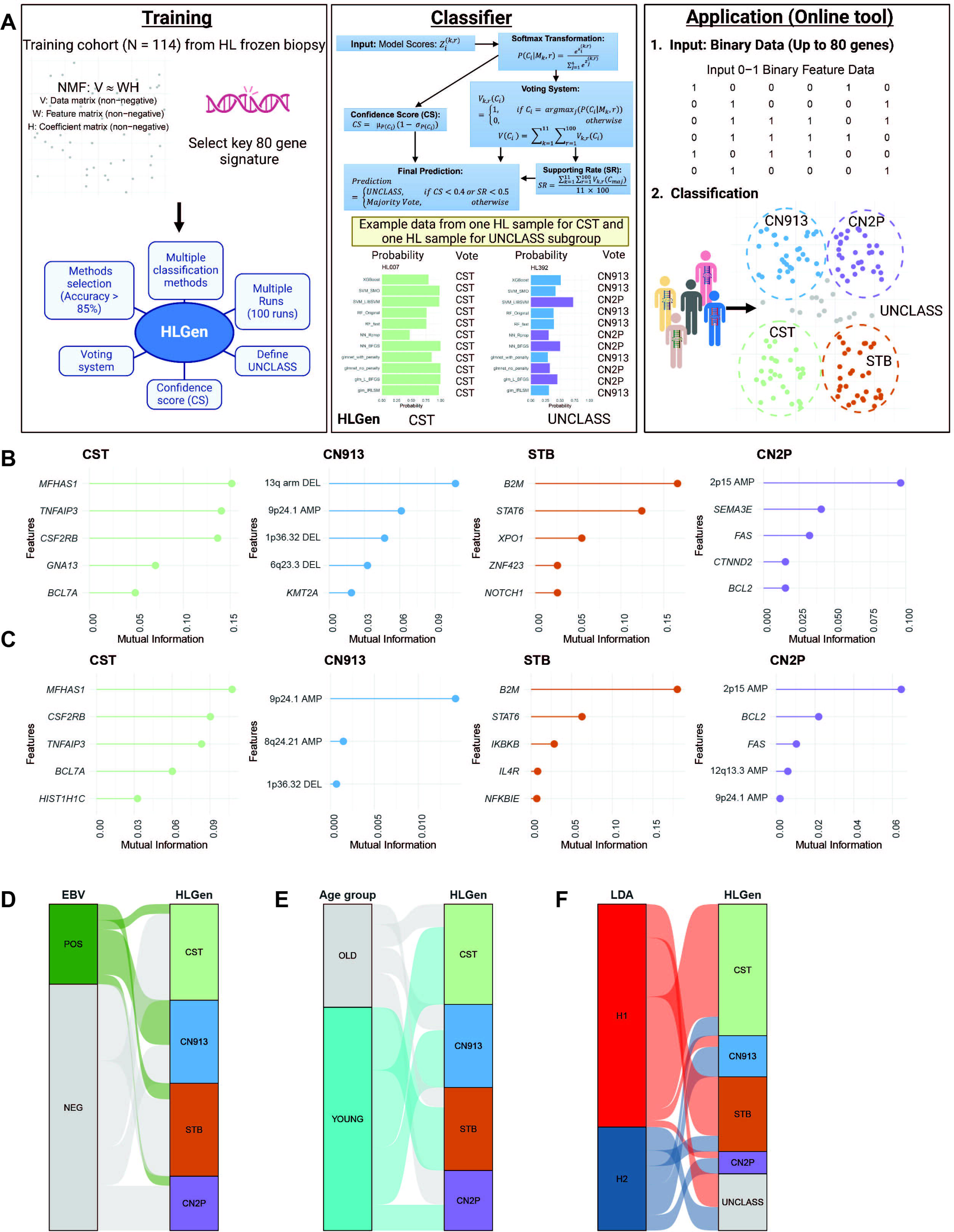
Establishment of HLGen molecular classification system. (A) Graphical summary of HLGen establishment including training process, classifier workflow, and development of an online tool. Training (left panel): The training pipeline starts with NMF clustering applied to 80 key genomic features from 114 HL frozen biopsy samples, shown in the top section. The bottom section summarizes the HLGen classifier, which integrates multiple classification methods, selects methods with > 85% accuracy, supports multiple runs, uses a voting system, calculates confidence scores, and defines the UNCLASS category for unclassifiable samples. Classifier (middle panel): This panel illustrates the HLGen classifier workflow, beginning with model scores as input, followed by a softmax transformation, the calculation of confidence scores, and the application of a voting system. The final prediction is determined by a combination of confidence scores, majority voting, and the supporting rate. The classifier is illustrated for n = 100 runs. The bottom section provides example outputs for two HL samples: one classified as “CST” and the other as “UNCLASS”, with bar plots showing the mean probabilities across all runs for each of the 11 selected classification methods. Application (right panel): The HLGen online tool requires binary input data with features matching the system’s defined names, available in the example dataset. Input samples are classified into one of four predefined classes or as UNCLASS, as shown in the bottom section. Figures showing the top five mutations or copy number alterations which contribute to each molecular cluster subtype in (B) Training cohort and (C) Stanford validation cohort. Figures showing the mutation or copy number alterations which contribute to each molecular cluster subtype in the Stanford cohort. The alluvial plot shows the association between the molecular subtypes defined by “HLGen” in the Stanford validation cohort(Alig *et al*., 2023) with (D) EBV status on HRS cells, (E) Age, and (F) LDA clusters as assigned by Alig et al.

## DISCUSSION

Our multidimensional tissue profiling revealed biological heterogeneity within CHL reflected by unique genetic profiles of malignant cells that are linked to TME composition and function **(Figure S18)**. Our multimodal profiling analysis provides an integrated view of molecular CHL subtypes that can be captured by HLGen, a novel molecular aberration-based classification tool. Recently, other groups have proposed molecular classification systems of CHL using ctDNA-based approaches(Alig *et al*., 2023; Heger *et al*., 2024). All proposed systems are concordant in their ability to identify a common group of tumors characterized by high mutational burden and CHL-specific copy number events. The HRS enrichment approach described here using fresh frozen tissue allowed not only to recover and validate previously proposed genotype clusters (H1/H2)(Alig *et al*., 2023), but also to increase discriminative power to further identify biologically distinct subgroups within simple mutation- and copy number-driven clusters.

As an example of the linkage between HRS cell mutational patterns and immune cells in evolving cellular ecosystems, we identified the biological mechanisms underlying immune regulatory TME milieus, driven by *CSF2RB* gain-of-function mutations. We identified novel and recurrent heterozygous truncating mutations in the cytoplasmic domain of CSF2RB leading to recruitment of Tregs *in vivo*. Notably, *CSF2RB* mutations are virtually absent in other major B-cell lymphoma entities, such as diffuse large B cell lymphoma (DLBCL) or follicular lymphoma(Pastore *et al*., 2015; Reddy *et al*., 2017; Schmitz *et al*., 2018; Wienand *et al*., 2019). Previous HL genomic studies have demonstrated the importance of the STAT6 signaling pathway in CHL pathogenesis(Alig *et al*., 2023; Tiacci *et al*., 2018; Weniger and Kuppers, 2021). In contrast, we here identified a STAT5-driven pathogenic mechanism in a subgroup of CHL. Of clinical importance, we demonstrated that selective JAK2 inhibition reversed the gain-of-function phenotype of *CSF2RB* mutations, indicating the potential benefit of targeting this pathway in the CST molecular subtype that encompasses tumors with *CSF2RB* mutations. This potential subtype-specificity of JAK2 inhibition might in part also explain the only marginal efficacy of JAK2 inhibitors observed in unselected and molecularly uncharacterized CHL patients in clinical trials(Van Den Neste et al., 2018).

CHL is currently classified into subtypes solely based on pathological features(Connors *et al*., 2020), while biological features are not incorporated in the existing pathological classification systems. The molecular classification system developed in this study, named HLGen, further improves biology-informed disease taxonomies, with the goal to guide treatment strategies and inform prognosis. Although purely based on genetic features, HLGen captures subgroups with distinct clinical features, such as a young age at diagnosis, and EBV-positivity of HRS cells. HLGen is publicly available as an online tool (https://shiny.bcgsc.ca/HLGen/), and requires only binary input data of up to 80 gene features. This approach offers flexibility in the number of gene features needed and the sequencing format as demonstrated in our ctDNA-based validation cohort. The treatment landscape of CHL has recently changed and is still evolving rapidly(Ansell et al., 2022; Castellino et al., 2022; Herrera et al., 2024). The impact of molecular profiles on clinical outcomes in the context of novel treatments, such as immunotherapy, therefore, needs to be elucidated in future studies. Notably, the HLGen molecular subgroups are each associated with specific immune cell populations expressing distinct co-inhibitory receptors, such as PD-L1, PD-1, and LAG3, indicating the potential of our molecular classification system as predictive biomarker framework for therapeutic target selection.

## METHODS

### Summary

Biopsies were selected from the biobank of BC Cancer’s Centre for Lymphoid Cancer based on availability of fresh frozen tissue samples and cell suspensions that were left over from routine diagnostic procedures. All cases were re-reviewed by experienced hematopathologists to confirm the diagnosis of CHL. We performed targeted capture sequencing and whole exome sequencing (WES) on HRS cells from fresh-frozen biopsies (n=114) obtained from our in-house lymphoma tumor bank at BC Cancer between 1989 and 2019 (“Discovery/Training Cohort”). HRS cells were enriched using laser microdissection (LMD) (n=108) or Flow cytometry (n=6) according to the availability of sample type, fresh-frozen tissue (FFT) or fresh frozen cell suspension, respectively, and DNA was extracted. Constitutional DNA was obtained from peripheral blood of 32 patients. Targeted sequencing was applied using a previously designed capture panel covering the coding sequence of 217 genes, as well as UTRs for 9 out of these 217 genes. Whole exome sequencing (WES) was applied to 6 tumor-normal pairs **(Table S2)**(Sarkozy *et al*., 2021). To identify potential outcome differences according to molecular features, univariate cox regression analyses were performed using data from pretreatment biopsies of CHL patients (n=103) who uniformly received first-line treatment with doxorubicin, bleomycin, vinblastine, and dacarbazine (ABVD) or ABVD-equivalent therapy with curative intent. In addition, we constructed tissue microarrays (TMA) from this cohort on which we performed GeoMx® Whole Transcriptome Assay (WTA) to obtain gene expression profiles from CD30+ HRS cells and multiplexed imaging analyses using multi-colour immunofluorescence (mIF) and imaging mass cytometry (IMC) to delineate the spatial TME ecosystem and its correlation with molecular feature of HRS cells(Chan et al., 2017). We utilized published sequencing data derived from circulating tumor DNA (ctDNA) (Stanford cohort)(Alig *et al*., 2023) to confirm clinical correlates. Patient characteristics of discovery cohort are summarized in **Table S1 and S3.**

This study was reviewed and approved by the University of British Columbia-BC Cancer Agency Research Ethics Board (H14-02304), in accordance with the Declaration of Helsinki.

### Laser microdissection (LMD)

LMD was performed as previously described using a Zeiss Axioplan 2 microscope equipped with Molecular Machines Industries technology in order to enrich for tumor cells(Sarkozy *et al*., 2021). Briefly, 6 µm FFT sections were fixed after cutting in 70% ethanol and mounted onto membrane slides. Membrane slides were stained for hematoxylin for 45 seconds, eosin for 15 seconds, dehydrated in ethanol bath (70%, 90% and 100%, 30 seconds each), incubated in Xylene for 1 minute and air dried. In order to dissect tumor cells, at least 10 sections per case were stained. Depending on the tumor content, tumor cells were individually picked or dissected in larger cell groups. For each case, at least 1200 tumor cells were collected. DNA extraction was performed using the DNA advanced Agencourt® extraction kit with overnight lysis. All DNA was then proceeded with library construction, without prior genome amplification.

### Sample preparation for flow cytometry-based cell sorting

To purify HRS cells, we implemented a flow cytometry-based cell sorting approach as described before(Aoki *et al*., 2024). In brief, cell suspensions from CHL tumors were rapidly defrosted at 37°C, washed in RPMI1640/20% FBS solution containing DNase I (Millipore Sigma, Darmstadt, Germany) and washed in PBS containing 2% FBS. Cells were resuspended in PBC containing 2% FBS and stained with antibody panel specific to isolate HRS cells for 15 minutes at 4°C in the dark. Viable cells (DAPI negative) were sorted on a FACS ARIAIII or FACS Fusion (BD Biosciences) using a 130 µm nozzle and were analyzed using FlowJo software (v10.2; TreeStar, Ashland, OR, USA). Sorted cells were collected in 0.3 ml of medium, centrifuged and diluted in 1x PBS with 0.04% bovine serum albumin (BSA). Cell number was determined using a Countess II Automated Cell Counter whenever possible.

### Capture design and sequencing

A detailed description of the targeted capture design and technical procedures can be found in *Sarkozy at al, Blood 2021*. Consensus variant calling was performed using VarScan (*version 2.4.3-0*) and Strelka (version 2.9.9.0)(Koboldt et al., 2009; Saunders et al., 2012),with single nucleotide variants (SNVs) requiring support from both variant callers, and insertions or deletions called by Strelka only. Paired somatic variant calling was performed on tumor samples for which paired constitutional blood DNA was available (n=26). The analysis was restricted to nonsynonmous variants. In addition, synonymous variants affecting *SOCS1*, *BCL7A* and *NFKBIZ* and located in regions that are putative targets of aberrant somatic hypermutation (aSHM) **(Table S2-3)** were retained. Somatic variants were identified in 106 genes **(Table S3)**. For tumor samples for which no paired normal sample was available (n=82) we used a pool of constitutional normal samples and restricted the analysis to the 106 genes identified in the paired analysis. To remove putative germline variants, we removed variants annotated in the gnomadAD database (https://gnomad.broadinstitute.org/) at a population frequency > 0.0001.

### Whole exome sequencing

Whole exome sequencing (WES) was performed on 6 tumor-normal pairs, following a previously established pipeline(Sarkozy *et al*., 2021). Analysis was restricted to the 106 genes identified to be somatic in the paired tumor-normal pairs that were analyzed using capture sequencing.

### Copy number calling

Copy number alterations (CNAs) were called from deduplicated bam files using CNVkit(Talevich et al., 2016). Bed files were created with the removal of inaccessible regions using a precomputed file for hg19, which excludes both the UCSC/ENCODE Duke mappability regions and the DAC blacklisted regions. Bin sizes for on- and off-target regions were estimated based on the median BAM file size using the “autobin” function. To enhance performance, all germline samples were combined to create a pooled reference. The hg19 reference genome was used to estimate GC content and repeat-masked proportions of each region. To identify relevant regions of CNAs, the GISTIC (Genomic Identification of Significant Targets in Cancer) method was applied to the curated segmentation results(Mermel et al., 2011). The analysis was performed using the web interface (http://genepattern.broadinstitute.org) with a threshold of 0.3 and −0.4 to define amplifications and deletions, respectively, a maximum of 10,000 segments, a q-value threshold of 0.25, confidence level of 0.9 (used to calculate the region containing a driver), and a focal length cut-off of 0.50. For each significant region, a peak region was identified, which is defined as the part of an aberrant region with the greatest amplitude and frequency of alteration and a minimum q-value.

### NMF clustering

NMF was applied using the NMF R package (Lee algorithm, k=2-7, n=100, random seed) to a binary matrix representing the mutational status of 73 genes (somatic mutation=1, no somatic mutation=0) that were found to be mutated in at least 5 tumor samples (5.7%) and 7 recurrent copy number gain (gain =1, no gain = 0) or deletion (deletion = 1, no deletion =0) events for which published literature provides several lines of evidence to support their role in CHL development (gain of 2p15 (*REL*), 8q24.21 (*MYC*), 9p24.1 (*CD274*, *PDCD1LG2*, *JAK2*) and 12q13.3 (*STAT6*), loss of 13q (arm-level), 1p36.31 (*TNFRSF14*) and 6q23.3 (*TNFAIP3*)). The factorization rank k = 4 was selected based on the optimal combination of cophenetic score and its drop-off(Nabet et al., 2024)**(Figure S10)** and sufficient granularity (4 clusters) to provide biological insight.

### Molecular classification development

We implemented an ensemble learning approach that integrates 16 statistical and machine learning classification algorithms. These algorithms were selected to include different models, architectures, optimization functions, and optimizers. Methods that achieved an accuracy higher than 85% (13 out of 16, 88%) in our training cohort were chosen. Among 13 methods, two methods would not fit to our online platform environment because of dependency to the “brulee” R package and removed. As a results, 11 methods were used to construct our final classifier (“HLGen”).

### Application of the HLGen classifier to an external cohort

Our newly developed HLGen classifier was applied to targeted sequencing and copy number calls from a recently published CHL study cohort by Alig et al (validation cohort)(Alig *et al*., 2023).The HLGen classifier incorporates the binary somatic status of 73 genes, 44 of which (60.3%) were covered by the targeted capture panel used by Alig et al(Alig *et al*., 2023). Notably, 38 out these were also part of Alig et al’s LDA-based classifier. Variant calls were obtained from https://hodgkin.stanford.edu(SNVs (CHL_Nature2023_SNVs_CAPPSeq.xlsx), and indels (CHL_Nature2023_Indels-CAPPSeq.xlsx)). Variant calls were filtered to include only the 44 overlapping HLGen genes and were restricted to nonsynonymous variants. In the case of *BCL7A* and *SOCS1*, synonymous variants affecting regions known to be targets of aSHM **(Table S3)** were retained. Cytoband-level copy number levels, obtained through their in-house pipeline, were kindly provided by *Dr. Ash Alizadeh*. CN calls were restricted to the cytobands that are interrogated by the HLGen classifier, and only high confidence CNA calls were retained by applying a FDR cut-off of 0.5%. Filtered variant calls and copy number status where then converted into a binary matrix, as described in the paragraph on NMF clustering. Clinicopathological annotations (including age and EBV status) for this cohort were provided by Dr. Ash Alizadeh.

To obtain stable cluster consensus calls from each HLGen algorithm they were each run 100 times on the Stanford data. An ensemble classification label was then obtained for each sample by applying majority voting and confidence score cutoffs across all runs and methods **(Figure 5A)**.

In our HLGen framework, ambiguous samples that do not clearly fit into any of the predefined classes are designated as “UNCLASS.” A sample is categorized as UNCLASS if either the proportion of supported votes is below 50% or the confidence score is below 0.4. The confidence score is mathematically defined as:

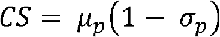

where *µ_p_* and *σ_p_* represent the mean and standard deviation of the classification probabilities, respectively. These values are estimated based on the observed classification probabilities across different methods and runs.

### Tissue microarray (TMA) construction, single color IHC and EBER-1 in situ hybridization (ISH) on TMA

For TMA construction, 1.5 mm duplicate cores were obtained from representative areas containing Hodgkin Reed-Sternberg cells of 114 biopsies of CHL. The diagnosis was made according to the WHO classification(Swerdlow *et al*., 2016). For IHC staining, 4 µm slides of the TMA and antibodies listed in Table S4 were used. Staining was performed on a Benchmark XT platform (Roche Diagnostics, USA) or intelliPATH platform (Biocare Medical, USA). The slides were independently scored by KT and/or TT and/or PF. Epstein-Barr virus-encoded small RNA 1 (EBER-1) ISH was performed according to the manufacturer’s protocol (Roche Diagnostics, USA). Detailed antibody information is listed in **Table S13.**

### Survival analysis

Association between the mutational status of recurrently mutated genes and clinical outcome were assessed by univariate CoxPH regression, using Progression Free Survival (PFS), defined as progression, relapse or death from any cause, as endpoint. The Kaplan-Meier method was used to estimate the survival functions, with log-rank tests performed to compare groups (*R package survival, version 3.3.1*).

### Spatial transcriptomics

For spatial transcriptomic analysis, we applied GeoMx Whole Transcriptome Assay (WTA) (version 1.0) on the tissue microarrays (TMA) of the exome/targeted sequencing cohort. Sections were cut at 5 μm thickness and mounted on plus-charged slidesthen processed on the BOND RX according to Semi-Automated slide preparation protocol (Leica Biosystem). Briefly, after baking the slides for 30 minutes at 60°C, slides were washed and incubated in the BOND RX with 1 µg/mL proteinase K. Next, slides were washed with PBS and incubated overnight at 37°C in an oven with 200 µl hybridization solution. Following hybridization slides were washed with stringent wash solution. Slides were incubated for 30 minutes at room temperature in a humidity chamber with Buffer W (NanoString) followed by morphology marker staining for 1 hour at room temperature with 200 µl morphology marker solution. Following staining, slides were scanned to capture fluorescent images used to select regions of interest (ROIs). CD30 and CD3 were used as morphological markers to identify and select CD30-positive regions (HRS-cells) from 95 patients as ROIs **(Figure S19)**. From 5 out of 95 patients, CD30-negative regions were selected as a negative control. Probe identities were captured via UV illumination and were transferred to 96 well PCR plates. Library preparation was performed with pro Code primers according to NanoString GeoMx DSP user manual. After library pooling and product purification, samples were sequenced on an Illumina Novaseq 6000 instrument. GeoMx data were further processed through GeoMx NGS pipeline and analyzed using the R package GeomxTools, version 3.1.1. Segments that did not meet the QC-cut-offs (Minimum number of reads per segment: 1000, minimum % of reads trimmed = 80%, minimum % of reads stitched = 80%, minimum % of reads aligned: 75%, minimum sequencing saturation: 50%, minimum negative control counts = 2, Maximum counts observed in NTC well = 1000, minNuclei = 25, minimum segment area = 5000) or the limit of quantification cut-off (2% of genes) were excluded from further analysis. After filtering, 44 CD30-positive and 5 CD30-negative high-quality segments were retained, and counts were normalized using full quantile normalization. DGEA was performed using a linear mixed model (LMM) to detect significantly differentially expressed genes between CD30-positive and CD30-negative regions. Preranked gene-set enrichment analysis (GSEA) was performed using the product of the log10(*p*-value) and direction of fold change for gene ordering (*R package fgsea, version 1.20.0*). A previously established HRS gene expression signature(Steidl *et al*., 2012) was then used to confirm HRS-specific expression in CD30-positive segments. Single sample gene set enrichment analysis (ssGSEA) was applied to calculate the degree of up-or downregulation for each pathway within each sample (*R package GSVA, version 1.42.0*). These ssGSEA enrichment scores were then subjected to Wilcoxon signed rank tests to identify pathways that were overexpressed in the CHL molecular subtypes or correlated with the mutational status of genes of interest, or MHCI/II expression on HRS cells.

### Imaging mass cytometry (IMC)

IMC was performed on a 5μm section of the TMA as described previously(Aoki *et al*., 2024) using a customized marker panel for CHL **(Table S4 and Figure S11)**. In brief, slides were imaged using the Fluidigm Hyperion IMC system with a 1 µm laser ablation spot size and frequency of 100-200 Hz. Tissue areas of approximately 1000 µm^2^ per sample were ablated and imaged. Segmentation was performed by the Ilastik machine learning package(Berg et al., 2019) in conjunction with the Python IMC Segmentation Pipeline published by the Bodenmiller group(Jackson et al., 2020). Cell type phenotyping was performed using a meta-clustering strategy. First, unsupervised clustering was performed on the transformed matrix of each ROI with first-tier cell phenotyping markers (CD3, CD4, CD8a, CD11b, CD14, CD20, CD30, CD68, CD123, Vimentin, Fibronectin, FOXP3) using Rphenograph (v0.99.1) with k = 15. Meta-clustering was then applied to the mean expression of these clusters with k=15, resulting in clusters categorized as T cells, B cells, myeloid cells, stromal cells, and HRS cells. Second, unsupervised clustering was performed individually on each cell subset (as defined by first cluster assignment) using cell-type specific markers to further type immune subsets **(Table S14)**: T-cell markers (CD4, CD8a, CD39, CD45RO, CCR4, CXCR3, CXCR4, CXCR5, ICOS, CXCL13, LAG3, TIM3, FOXP3, CTLA4, PD1, GranzymeB, BCL6, and Ki67), B-cell markers (CD20, CD27, CD11c, CXCR5, BCL6, and Ki67), myeloid markers (CD4, CD11b, CD11c, CD14, CD68, CD80, CD123, CD163, Galectin9, PDL1, TIM3, GranzymeB, CXCL13, and Ki67), stromal-cell markers (aSMA, Vimentin, Fibronectin, CXCL12, CXCL13, and Ki67), and HRS-cell markers (TARC, CD30, CD47, CD80, CD123, CXCR4, CXCR5, CLEC2D, Galectin9, TGFβ, PDL1, and Ki67). The following criteria were used: Markers used to type immune subsets included CD3, CD8, GranzymeB (cytotoxic T cells, CTL), PDCD1, CXCR5, BCL6 (TFH cells).

For each cell, we calculated the spatial interaction score, called ‘spatial score’, to a given cell type as the distance to the 5 nearest neighbor cells, capped at the spatial interaction range, scaled, and inverted as described previously(Aoki *et al*., 2024). The CellChat package (v2.1.0) was used to identify potential cell-cell communication networks, incorporating both protein expression and spatial information. Ligand-receptor pairs were obtained from CellChat’s built-in databases, which included antibodies present in the IMC panel. Communication probability was calculated for each ligand-receptor pair, and significant interactions were identified at a threshold of *p*<0.05.

### Cell lines

Human CHL-derived cell lines L-428, and U-HO1 were purchased from DSMZ (German Collection of Microorganisms and Cell Cultures GmbH; Braunschweig, Germany). The 293FT cells were purchased from Invitrogen (catalog no. R70007). The L-428 cells were cultured in Roswell Park Memorial Institute-1640 medium (RPMI-1640) supplemented with 10% FBS (Corning, catalog no. 35-077-CV), and U-HO1 cells were cultured in RPMI-1640 and Iscove’s Modified Dulbecco’s Medium (IMDM) (1:4) supplemented with 20% FBS. The 293FT cells were cultured in in Dulbecco’s modified Eagle’s medium (DMEM) supplemented with 10% FBS. FBS was heat-inactivated by heating serum at 56°C for 30 minutes before addition to the medium. All cell lines were authenticated by short tandem repeats (STR) profiling (The Centre for Applied Genomics, Toronto, Canada) and confirmed to be negative for mycoplasma contamination using the Venor GenM Mycoplasma Detection kit, PCR-based (Sigma Aldrich, catalog no. MP0025). For the stimulation experiments, recombinant human IL-3 (1ng/mL, PeproTech, catalog no. 200-03), recombinant human IL-5 (1ng/mL, PeproTech, catalog no. 200-05), recombinant human GM-CSF (10ng/mL, PeproTech, catalog no. 300-03) were used.

### CSF2RB site-directed mutagenesis and viral transduction

The wild-type (WT) CSF2RB coding sequence was amplified by polymerase chain reaction (PCR) using complementary DNA from U-2940 cells and cloned into the pRETRO-delTO-CMV-Puro-mCherry retroviral plasmid vector (pRETRO). The S723*, E788*, K838*, Q853*, Q867*, and Q885* mutations were introduced using the GENEART site-directed mutagenesis system (Thermo Fisher Scientific, catalog no. A14604) according to the manufacturer’s instructions. Viral particles were generated in 293FT cells and concentrated using the Lenti-X concentrator (Takara Bio, catalog no. 631231). L-428 and U-HO1 isogenic cells were sequentially engineered with: 1) feline endogenous virus expressing the ecotropic retroviral receptor, and 2) retrovirus containing the pRETRO-CSF2RB-WT and mutants. Retroviral transduction of these cells, and mCherry-positive fraction cell sorting followed by puromycin selection (4 µg/mL; InvivoGen, catalog no. ANT-PR-1) was performed as previously described(Alig *et al*., 2023; Vigano et al., 2018).

### Flow cytometry

Flow cytometry for surface CSF2RB (CD131) and intracellular STAT5 phospho-Y694 was performed on a BD FACSymphony™ Flow Cytometer (Becton-Dickinson Biosciences) using the following primary antibodies: BD OptiBuild™ BV421 Mouse Anti-Human CD131 (3D7), BD Horizon™ BV421 Mouse IgG1, k Isotype Control(X40), and Phosphor-STAT5 (Y694) Rabbit mAb (C11C5) (all dilution, 1:200). For the intracellular staining, cells were fixed with 2% paraformaldehyde in PBS with 2% FBS for 15 minutes, and permeabilized with Cold MetOH for 30 minutes on ice. The cells were stained with primary antibody for 2 hours, followed by secondary staining using Goat anti-Rabbit IgG (H+L) Cross-Adsorbed Secondary Antibody, Alexa Fluor™ 647 (Invitrogen, catalog no. A-21244) for 30 minutes. Detailed Ab information is listed in **Table S15**.

### Immunoblotting

Immunoblotting was performed as previously described(Alig *et al*., 2023) using the following primary antibodies: STAT1 Rabbit mAb (D1K9Y), STAT3 Mouse mAb (124H6), STAT6 Rabbit mAb (D3H4), AKT(pan) Rabbit mAb (C67E7), Phospho-STAT1 (Y701) Rabbit mAb (58D6), Phospho-STAT3 (Y705) Rabbit mAb (D3A7), Phospho-STAT5 (Y694) Rabbit mAb (C11C5), Phospho-STAT6 (Y641) Rabbit mAb (C11A12), Phospho-AKT (S473) Rabbit mAb (193H12), GAPDH Rabbit mAb (14C10) (all from Cell Signaling Technology, dilution 1:1000), STAT5 Rabbit polyclonalAb (C-17) (dilution 1:5000), CSF2RB (IL-3/IL-5/GM-CSFRβ Mouse mAb (A-3) (dilution 1:100) (Santa Cruz Biotechnology), followed by secondary staining using anti-Mouse (dilution 1:10000; Promega, catalog no. W4021) or Rabbit IgG (H+L) HRP conjugate (dilution 1:5000; Promega, catalog no. W4011), and detected using Amersham ECL Detection Reagents (Cytiva, catalog no. RPN3004). Detailed Ab information is listed in **Table S16**.

### Human CCL17/TARC and IL-13 detection by ELISA, and IL-5 detection by Luminex assay

Enzyme linked immunosorbent assay (ELISA) was performed to quantitate CCL17/TARC and IL-13 secretion in the L-428 and U-HO1 cells. To measure cytokine and chemokine production, we harvested culture supernatant after 48 hours of culture from and assayed using the Human CCL17/TARC (Catalog #DY364-05), IL-13 (Catalog # DY213-05), and DuoSet ELISA Ancillary Reagent Kit 2 (R&D Systems, catalog no. DY008B) according to the manufacturer’s instructions. The levels of IL-5 were measured using a 35-plex Luminex assay kit (Thermo Fisher Scientific, catalog no. LHC6005M) according to the manufacturer’s protocol.

### Quantitative RT-PCR

Quantitative RT-PCR was performed as previously described(Vigano *et al*., 2018). TaqMan gene expression assay probes were used to detect messenger RNA (mRNA) levels using the following probes: Human CSF2RB (Assay ID: Hs001666144_m1), Human CCL17 (Assay ID: Hs00171074_m1), Human IL-13 (Assay ID: 00174379_m1) (Thermo Fisher Scientific), and Human GAPDH Endogenous Control (Applied Biosystems™, catalog no. 4333764F).

### RNA-seq

RNA-seq was performed on two biological replicates each from the L-428 (transduced with empty vector or a vector expressing CSF2RB-WT or CSF2RB-E778* and U-HO1 (CSF2RB-WT or CSF2RB-WT/E788*, stimulated with IL-3 or IL5) isogenic cell lines. Following quality control checks using a BioAnalyzer 2100 Instrument (Agilent, Santa Clara, California), 200 ng of RNA was subjected to library construction using the Illumina Stranded mRNA Prep Ligation Kit (Illumina, catalog no. 20040534), then run on an Illumina NextSeq550 using a High Output 150 cycle kit (Illumina, San Diego, California). This yielded on average 30 million paired-end reads per sample.

### RNA-seq analysis

RNA-seq reads were aligned to GRCh38 and per-gene counts were generated using the STAR aligner (version 2.5.3a_modified)(Dobin et al., 2013). Counts were normalized and scaled using trimmed mean of M-values (TMM) normalization and log2-transformed counts per million (CPM) scaling, respectively(Robinson et al., 2010). Differential gene expression analysis (DGEA) was performed using DESeq2 v1.26.0(Love et al., 2014). Gene Ontology (GO) analysis, interrogating a collection of MSigDB (Broad Institute) annotated gene-sets (Hallmark, BioCarta, KEGG, Reactome, and Gene Ontology (GO) terms, https://www.gsea-msigdb.org/gsea/msigdb), and signatures annotated in the LymphoChip database (https://lymphochip.nih.gov/signaturedb), was applied to identify enriched gene expression signatures among the most significantly differentially expressed genes(Yu et al., 2012).

### IL5RA neutralization and pacritinib inhibition experiments

L-428 cells were treated with Human IL-5 receptor alpha /CD125 Neutralization Antibody (10μg/ml, R&D Systems, catalog no. AF-253-NA) for 1 hour. For pacritinib inhibition experiments, pacritinib (Cayman, CAS No#937272-79-2) were added to L-428, and U-HO1 in the presence of IL-3 or IL-5 for 48 hours.

### T cell isolation and chemotaxis assays

CD8-CD3+ cells were purified from human peripheral blood mononuclear cells (PBMCs) obtained from four different healthy donors using magnetic-activated cell sorting (MACS, Miltenyi Biotec, Gaithersburg, MD). Subsequently, the cells were stained on ice with the following reagents: i) Live/Dead Fixable Green (Invitrogen, catalog no. L23101) for 20 minutes, ii) Human BD Fc Block (BD Biosciences, catalog no. 564220) for 15 minutes, and iii) anti-human CD4 (BD Biosciences, Franklin Lakes, NJ) and CCR4 (BioLegend, San Diego, CA) for 30 minutes. Fluorescence-activated cell sorting was performed to isolate CD4+CCR4+ and CD4+CCR4- cells after gating for the live cell population. Sorted cells were cultured overnight in a complete medium (RPMI-1640 supplemented with 10% FBS) at 37°C. The percentages of CD25+FOXP3+ cells in pre- and post-sorted populations were quantified via FCM using anti-human CD25 (BioLegend, San Diego, CA) and FOXP3 (BD Biosciences, Franklin Lakes, NJ) **(Figure S20)**. For intracellular staining, the FOXP3/Transcription Factor Staining Buffer Set (Invitrogen, catalog no. 00-05523-00) was employed for permeabilization. Detailed antibody information can be found in **Table S15**.

T cell chemotaxis assays were conducted using HTS Transwell 96-well plates with 5.0 µm pores (Corning, Corning, NY). L-428 supernatant was prepared by culturing 1 × 10L cells/mL for 8 hours, followed by dilution in complete medium at a 1:20 ratio. Briefly, 1 × 10L sorted T cells in 80 µL of complete medium were pre-incubated at 37 °C for 30 minutes with either 5.0 µM CCR4 antagonist (Cayman Chemical Company, catalog no. 21885) or DMSO as a vehicle control. Without washing, the cells were transferred to the upper insert of the transwell and positioned above the lower chamber, which contained 100 µL of L-428 supernatant. After 2 hours of incubation, the CellTiter-Glo assay (Promega, catalog no. G7571) was performed by mixing 5 µL of migrated cell suspension with 95 µL of complete medium. Luminescence was measured using a Spark10M instrument (Tecan, Männedorf, Switzerland) according to the manufacturer’s instructions. To determine the percentage of migrated cells, 1 × 10L T cells were seeded directly into one bottom well as an input control.

### Engraftment of cord blood cells and tumor cells

CD34+ human cord blood (CB) cells (Eaves Stem Cell assay) were rapidly thawed at 37°C and resuspended in 1:1 cold PBS (Gibco, catalog no. 10010023) and FBS (Corning). Cells were centrifuged at 300 g for 5 minutes, resuspended in cold PBS at 5×10^5^ cells/mL, and kept on ice until injection. 5×10^4^ cells in 100 µL cold PBS were intravenously engrafted into the tail vein of female CNW (NOD.Cg-Rag1nullIl2rgnull/SzJ-W41/41) mice between 7-10 weeks old using 28G insulin syringes (BD Bioscience). 8 weeks following CB engraftment, 1×10^6^ luciferase-labeled L- 428 cells in 100 µL PBS were intravenously engrafted into the tail vein.

### Bone marrow aspiration

Red blood cells were lysed using RBC lysis buffer (BioLegend, San Diego, CA, catalog no. 420301) following the manufacturer’s protocol. Cell suspensions were stained on ice with i) Live/Dead Fixable Green (Invitrogen, Waltham, MA) for 20 minutes, ii) followed by Human Fc Block (BD Bioscience, Franklin Lakes, NJ) for 15 minutes, and anti-human CD45 (BioLegend, San Diego, CA) and CD45 (BD Bioscience, Franklin Lakes, NJ) for 30 minutes. Detailed Ab information is listed in **Table S15**.

### Tumor burden monitoring

D-luciferin, potassium salt (Goldbio, catalog no. LUCK-100) was resuspended in PBS (Gibco) at 15 mg/mL and 200 uL of the mixture was administered intraperitoneally. 10 minutes after the administration, mice were imaged by in vivo imaging system (IVIS) ® Lumina™ S5 Imaging System (Revvity, Massachusetts, USA) to assess tumor burden.

### Organ harvest

Spleens were harvested from all mice at the time of sacrifice. Each spleen was bisected using a sterile blade, ensuring both halves included tumor-infiltrated regions. One half of the spleen was mechanically disaggregated to obtain cell suspensions for FCM analysis or cryopreservation. The other half was fixed in formaldehyde overnight and subsequently transferred to 70% ethanol for histological analysis.

### FCM analyses for the humanized mice experiments

Single cell suspensions were stained on ice with i) Live/Dead Fixable near-IR red (Invitrogen, Waltham, MA) for 20 minutes, ii) followed by human Fc block (BD Bioscience, Franklin Lakes, NJ), and human immunophenotyping panel (anti-human CD45 [BioLegend, San Diego, CA], CD3, CD4, CD8, CD19, CD64 [BD Bioscience, Franklin Lakes, NJ], and CD14 [Invitrogen, Waltham, MA].) or human Treg panel (anti-human CD45, CD25, CCR4 [BioLegend, San Diego, CA], CD3, CD4, CD8, and FOXP3 [BD Bioscience, Franklin Lakes, NJ]). FOXP3 staining was performed following manufacturer’s protocol (Invitrogen, Waltham, MA). L-428 was identified by dual YFP and mCherry expression after gating for live cells. Samples were collected using Symphony A5 (BD Bioscience, Franklin Lakes, NJ) and data were analyzed with FlowJo version 10.9.0 (FlowJo LLC, Ashland, OR). Detailed Ab information is listed in **Table S15**.

### Statistical results & visualization

Statistical analyses were performed with Prism software (GraphPad Prism 10.4.0), using unpaired two-sample *t*-test, paired two-sided t-test, one-way ANOVA, and negative binomial regression model, where appropriate. The Dunnett’s multiple comparisons test was carried out to control the false discovery rate for multiple comparisons. P-values < 0.05 were considered to be statistically significant. Asterisks represent associated p-values (\**p*<0.05; \*\**p*<0.01; \*\*\**p*<0.001; \*\*\*\**p*<0.0001). In all boxplots, boxes represent the interquartile range with a horizontal line indicating the median value. Whiskers extend to the farthest data point within a maximum of 1.5 × the interquartile range, and colored dots represent outliers. BioRender (BioRender.com) was used to create the schematic outlines.

## Data availability

The raw sequencing data described in this study will be deposited in European Genome-phenome Archive (EGA, accession numbers TBD) and will be available by request to the authors. Our molecular classification system, “HLGen” is available at https://shiny.bcgsc.ca/HLGen/.

## Code availability

Scripts used for data analysis are available upon request.

## AUTHORS’ CONTRIBUTIONS

Study design: T.A., S.R., A.M., and C.S.; Writing: All Authors; Manuscript review: All Authors; Data interpretation: T.A., G.D., S.R, A.J., M.K., M.L., L.H., S.A., M.S.E, A.A., and C.S.; *In vitro* experiments: T.A., S.R., M.K., M.L. and D.S; *In vivo* experiments: T.A., S.R., M.K., M.L., G.E. and L.G. Data analysis: T.A., G.D., S.R, A.J., M.K., M.L., L.H., S.A. and S.H; DNA sequencing processing: T.A., C.S., L.O.B; IHC work: K.T., T.M-T., K.M., C.S., T.G., C.D.V. and P.F.; Pathological review: K.T., T.M-T. and P.F.; Case identification: T.A., S.R., J.P.R, M.H., M.C., J.K., A.P. and E.K.; IMC work: T.A., A.L., Y.Y., A.T., S.W., E.L, A.X., A.M., and A.R.; Supervision: A.P.W., D.H., B.N., R.D.M., A.A., K.J.S., D.W.S., and C.S.

## ACKNOWLEDGMENTS

This study is supported by Program Project Grant funding from the Terry Fox Research Institute (Grant No.1061 and 1108), Large Scale Applied Research Project funding from Genome Canada (Grant No. 13124), Genome BC (Grant No. 271LYM) and CIHR (Grant No.GP1-155873, Grant No. 191809), the Canadian Cancer Society Research Institute (Grant No. 705288), a Foundation grant from CIHR (Grant No. 148393), the BC Cancer Foundation and the Paul G. Allen Frontiers Group (Distinguished Investigator award to C.S., Grant No.12829). T.A. was supported by a fellowship from the Japanese Society for The Promotion of Science and the Uehara Memorial Foundation. T.A. was supported by a fellowship from CIHR, the Lymphoma Research Foundation and the Uehara Memorial Foundation. T.A. received research funding support from The Kanae Foundation for the Promotion of Medical Science. T.A. is the recipient of a Lymphoma Research Foundation Lymphoma Scientific Research Mentoring Program Scholarship award. S.R. was supported by the Randy Gascoyne lymphoma fellowship, and by a physician scientist fellow award from the Leukemia & Lymphoma Society of Canada.

