## Supplemental Figures for "Multidimensional characterization of cellular ecosystems in Hodgkin lymphoma"

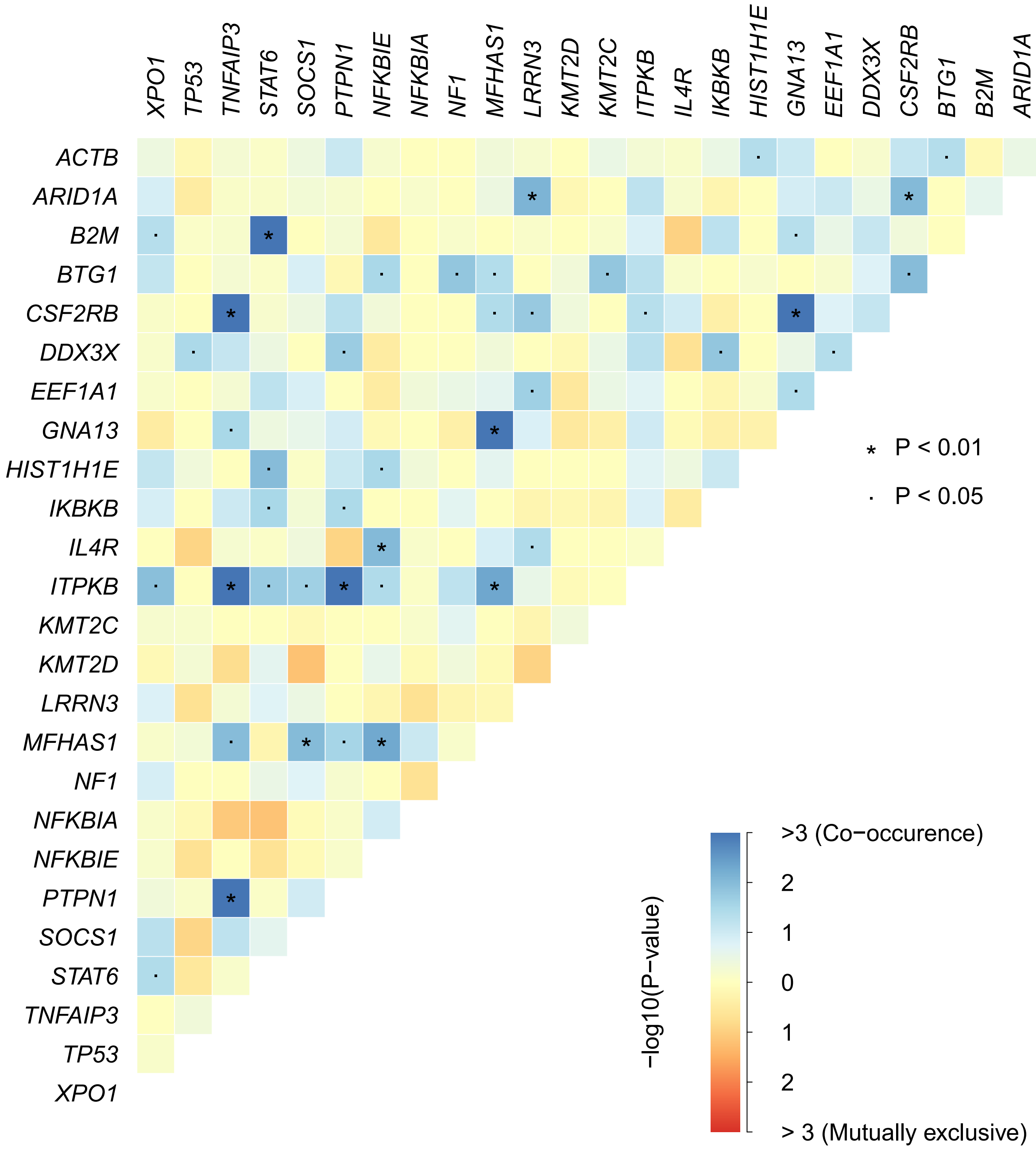


**Figure S1. Co-occurrence of mutation in CHL.** A mutual exclusivity/co-occurrence analysis was performed across 114 CHL samples.

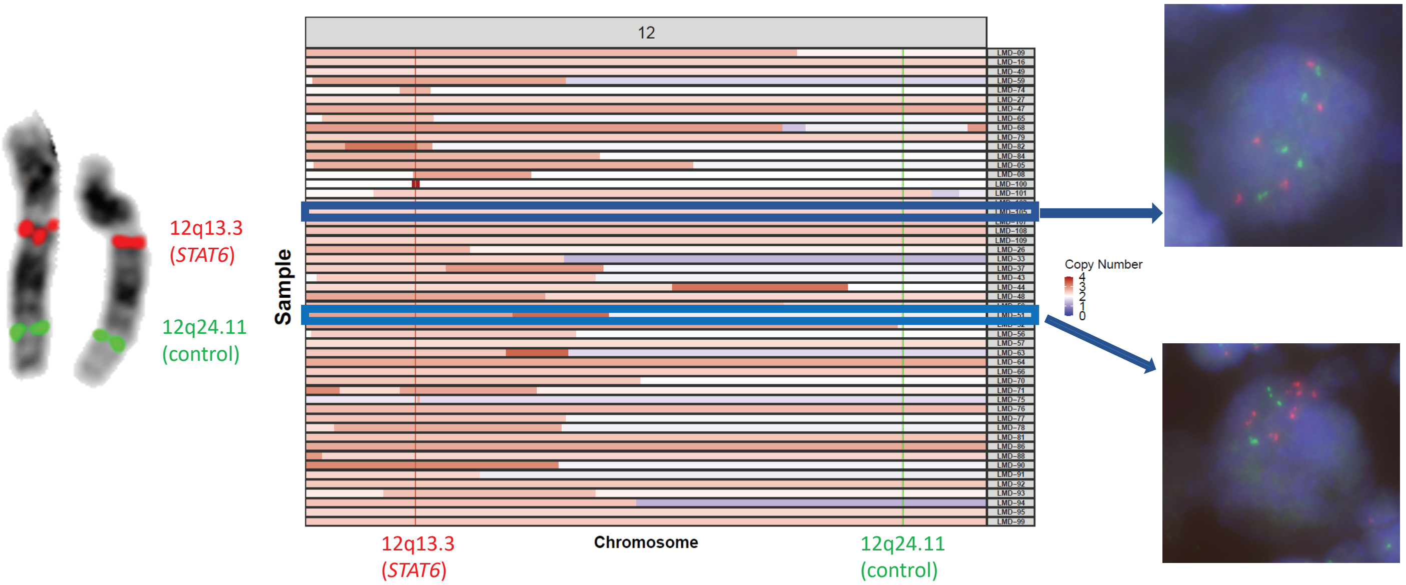


**Figure S2. *STAT6* copy number gain.** Copy number gains were validated using fluorescence in situ hybridization (FISH) assays. Two-color FISH assays use a green probe interrogating *STAT6* (chromosome 12q13.3) and a red probe interrogating chromosome 12q24.11 as a reference.

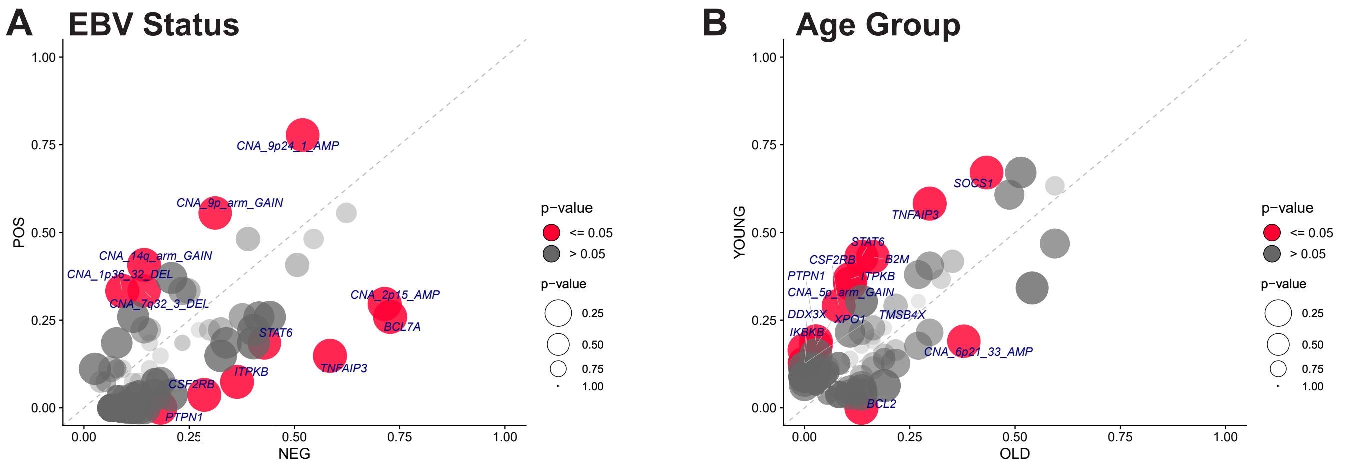


**Figure S3. Frequent mutations according to key clinical features.** Pairwise (Fisher's exact test) comparison of the incidence of mutations within **(A)** EBV + CHL vs EBV – CHL or **(B)** younger CHL (Age < 45) vs older CHL (Age >= 45). The size of the bubble is inversely correlated to the p-value.

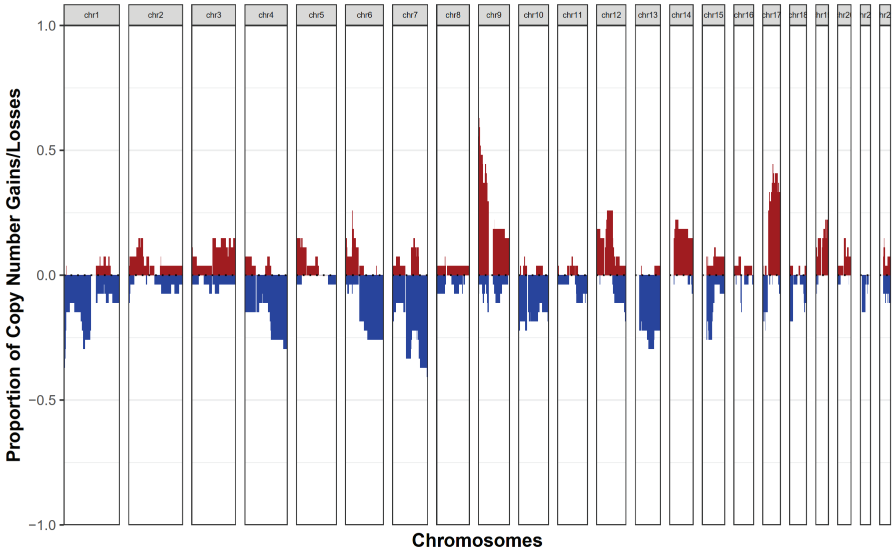


EBV +

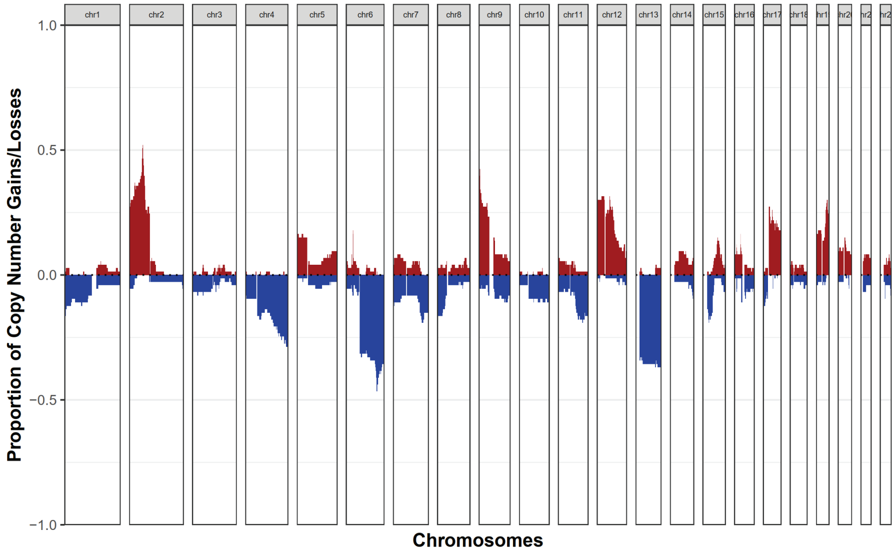


EBV -

**Figure S4. Copy number alterations according to EB Virus (EBV) status on HRS cells.** GISTIC plot highlighting the main copy number variation in CHL according to EBV status on HRS cells.

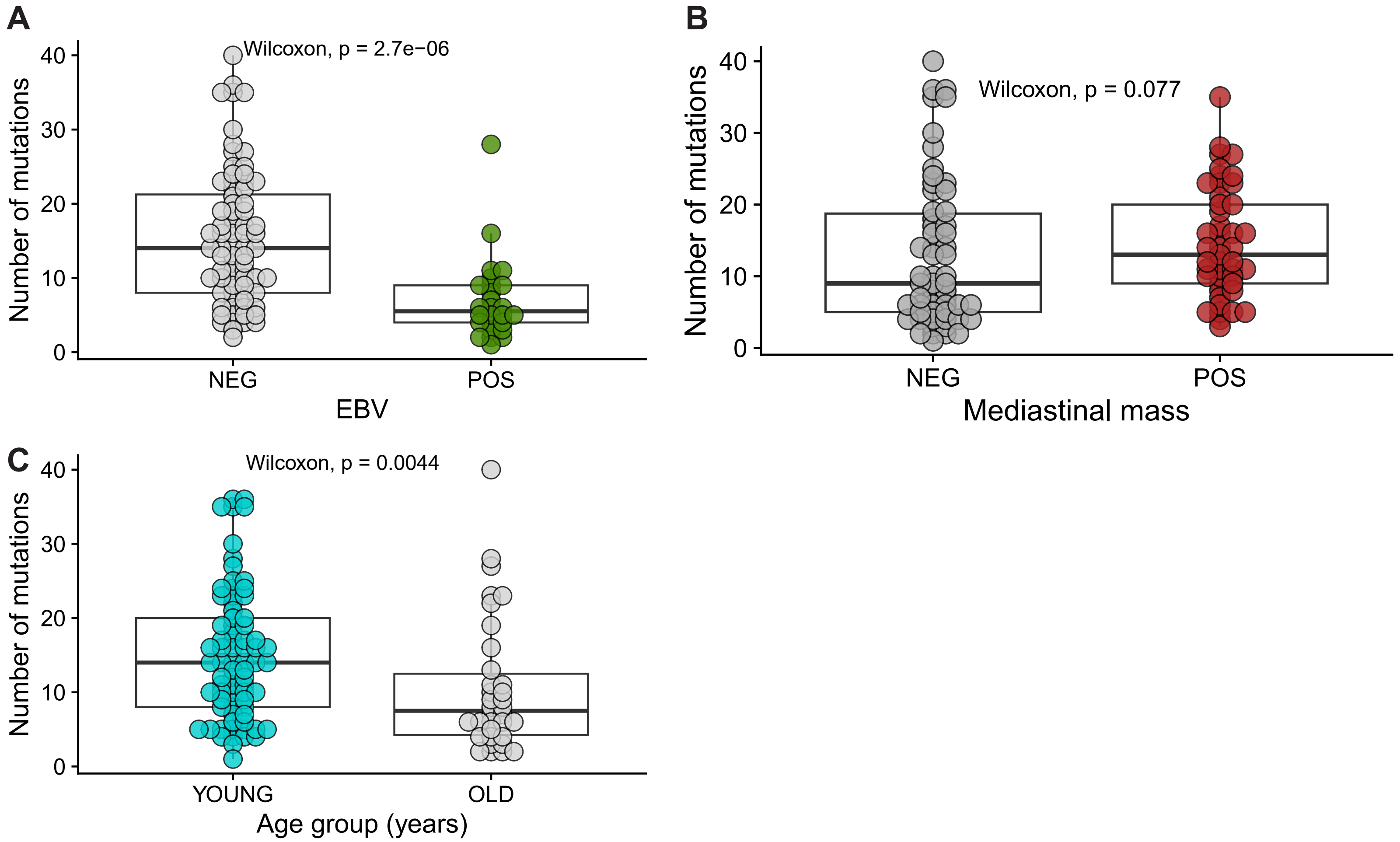


**Figure S5. Coding mutation count according to key clinical features in CHL.**
Coding mutation count within the 217 target genes according to **(A)** EBV status on HRS cells, **(B)** Mediastinal mass and **(C)** Age group.

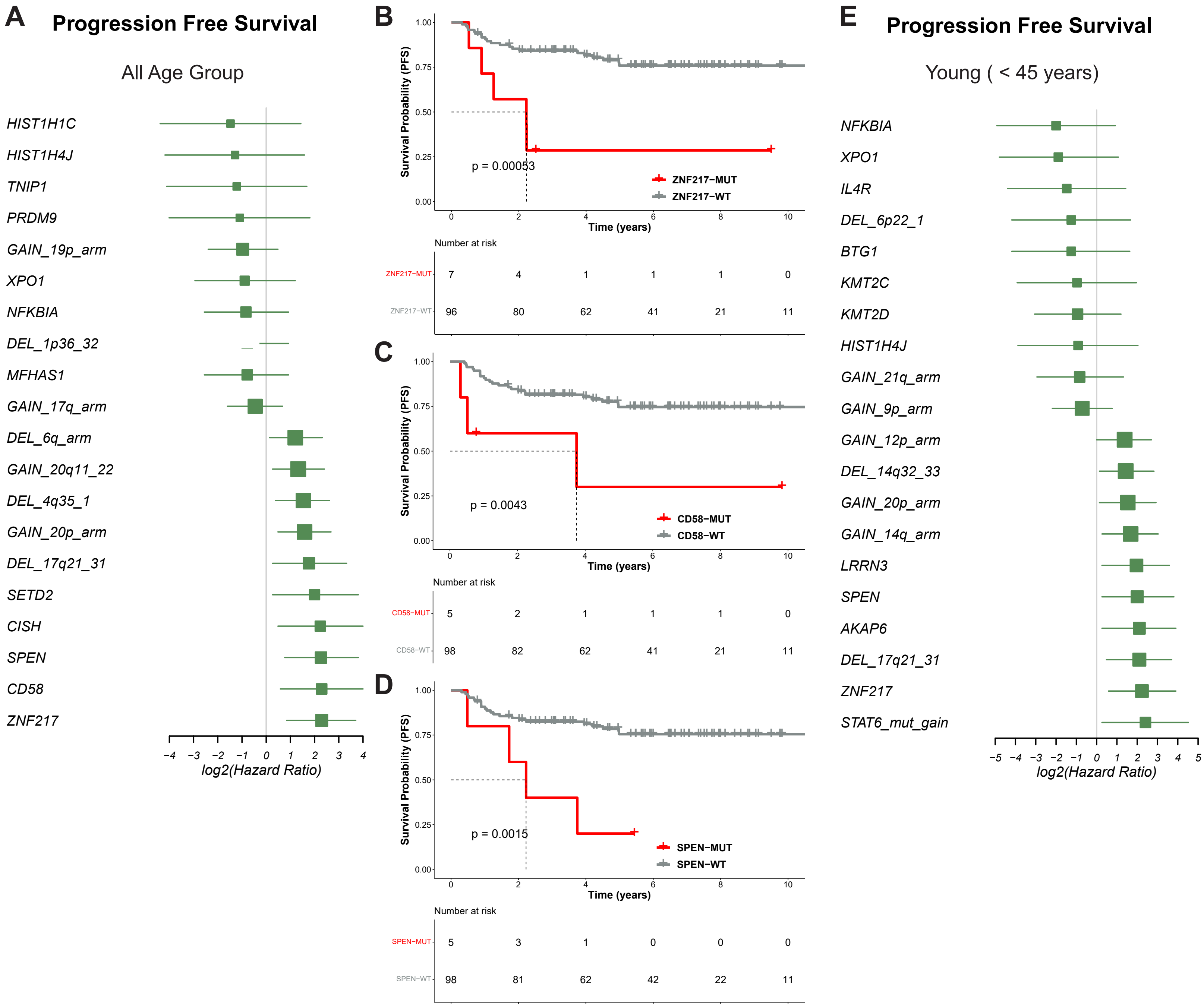


**Figure S6. Impact of molecular profile on survival in CHL. (A)** Forest plots summarize the mutation or copy number alteration associated with progression-free survival (PFS) in CHL uniformly treated with ABVD-like treatment (n=103). Kaplan-Meier curves for PFS according to mutational status in **(B)** *ZNF217*, **(C)** *CD58* and **(D)** *SPEN* for CHL patients. P values were calculated using a log rank test. **(E)** Forest plots summarize the mutation or copy number alteration associated with PFS in younger CHL (age < 45 years) uniformly treated with ABVD-like treatment (n=71).

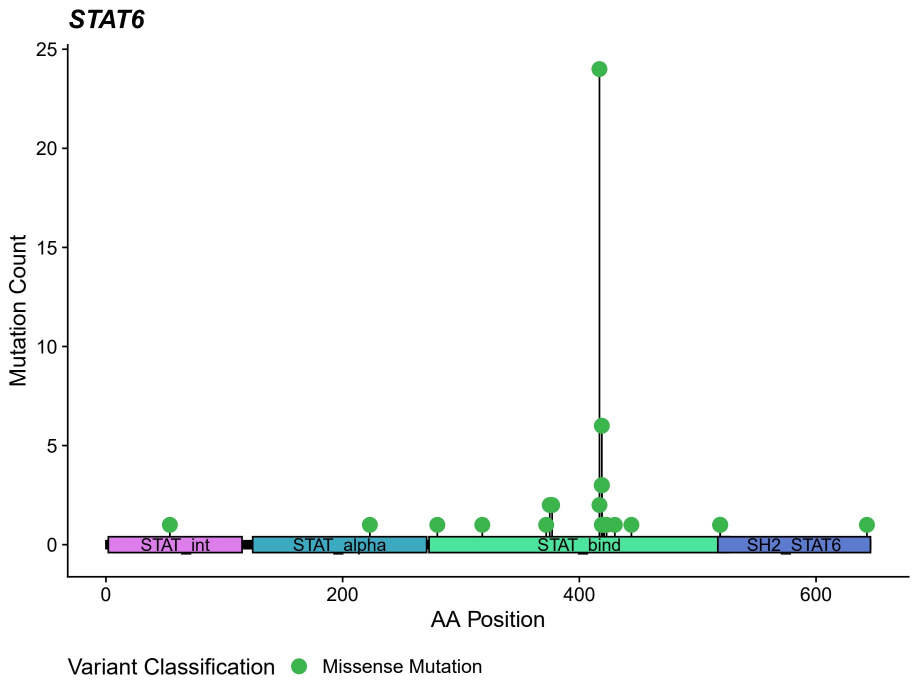


**Figure S7. *STAT6* mutations in CHL.** STAT6 domains protein and the locations of mutations identified in this study.

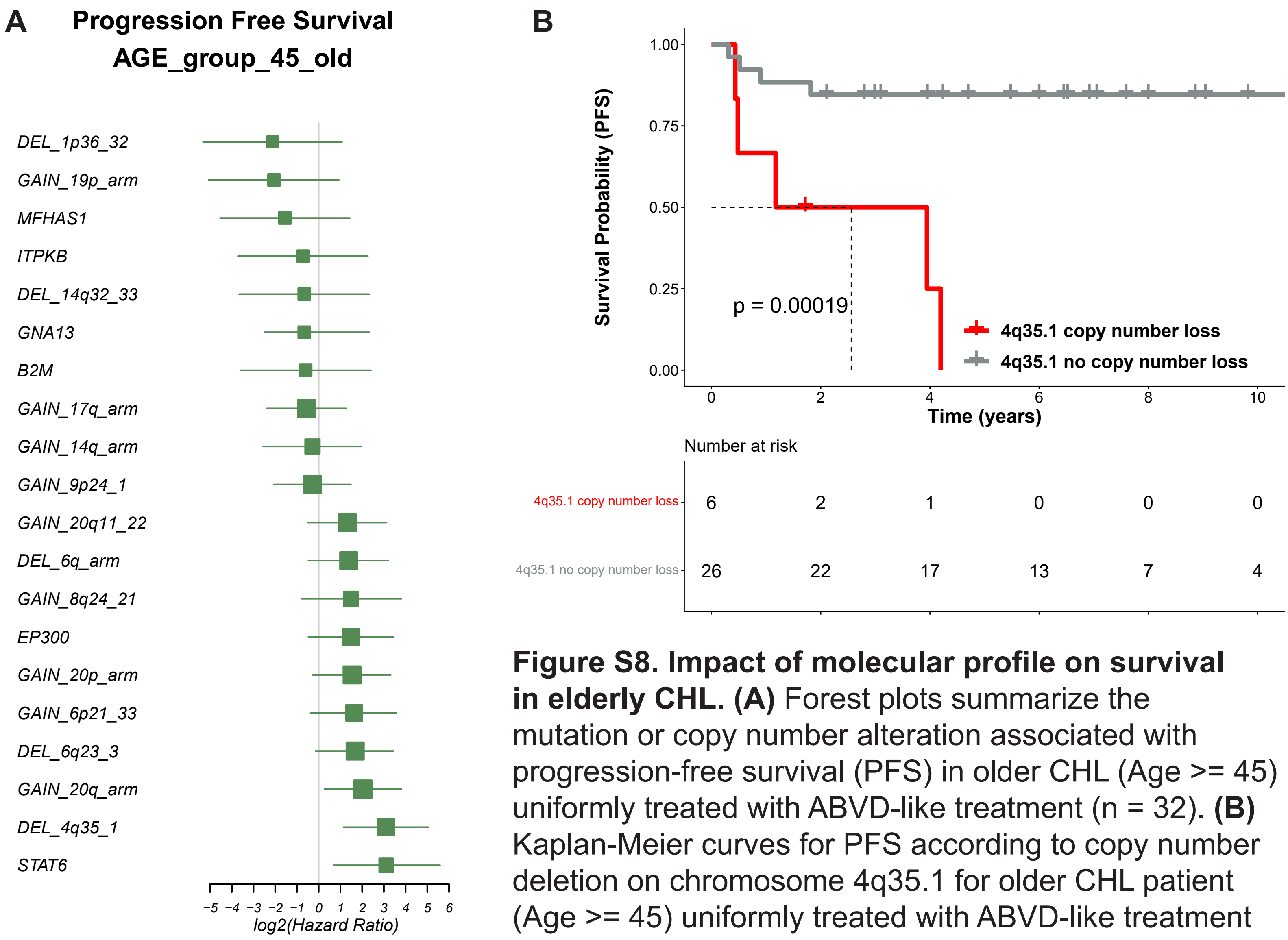


**Figure S8. Impact of molecular profile on survival in elderly CHL. (A)** Forest plots summarize the mutation or copy number alteration associated with progression-free survival (PFS) in older CHL (Age >= 45) uniformly treated with ABVD-like treatment (n = 32). **(B)** Kaplan-Meier curves for PFS according to copy number deletion on chromosome 4q35.1 for older CHL patient (Age >= 45) uniformly treated with ABVD-like treatment (n = 32). *P*-values were calculated using a log rank test.

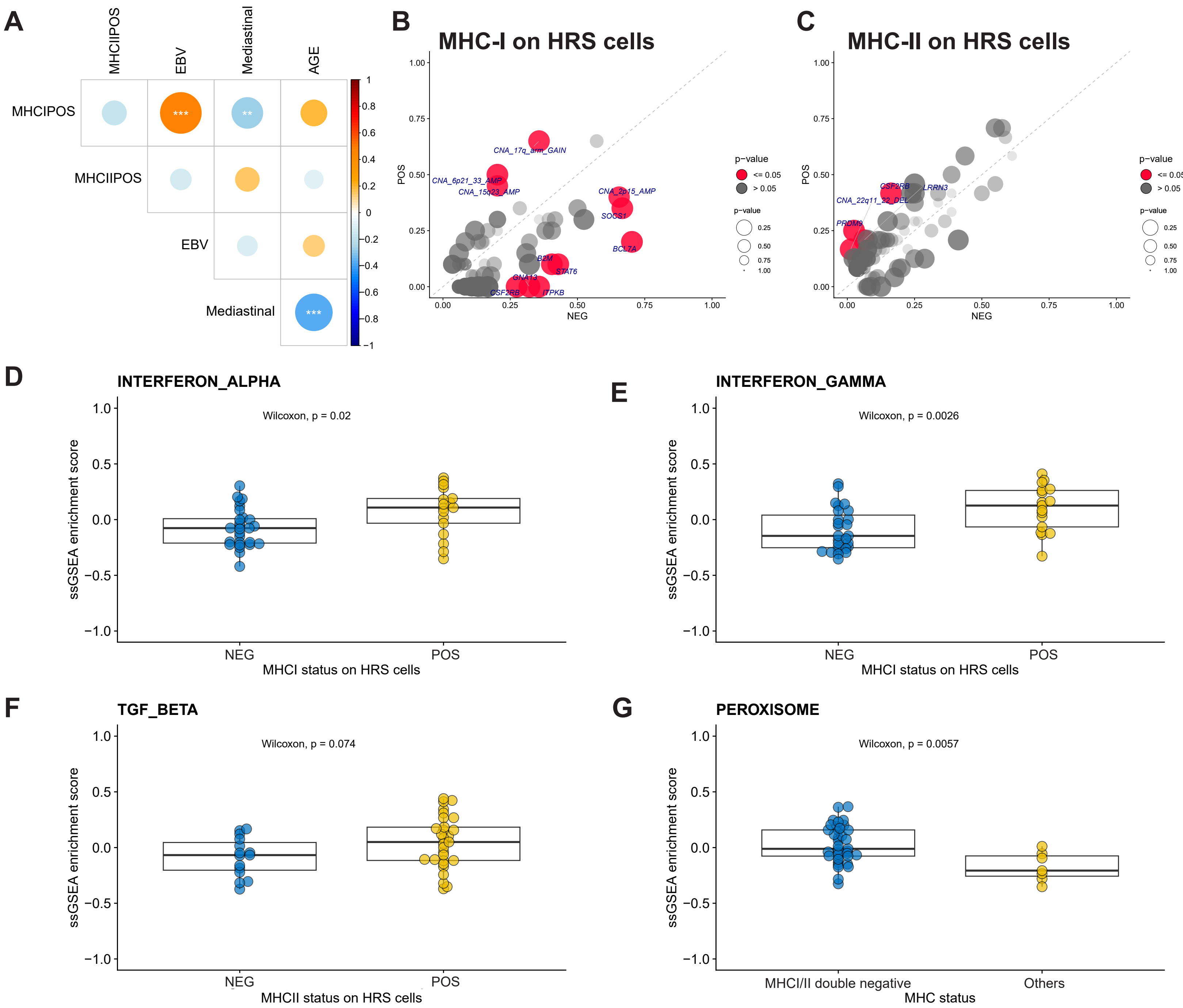


**Figure S9. MHC status on HRS cells and their molecular correlates. (A)** Dot plot showing correlations between MHC status on HRS cells and major clinical features. Dot size and color summarize Pearson correlation values, with positive correlations represented in red and negative correlations represented in blue. Asterisks represent associated p-values (**p*< 0.05; ***p*< 0.01; ****p*< 0.001). **(B-C)** Pairwise (Fisher's exact test) comparison of the incidence of mutations within **(B)** MHC Class I (MHC)+ CHL vs MHC-I -CHL or **(C)** MHCII+ CHL vs MHC-I - CHL. The size of the bubble is inversely correlated to the *p*-value. Box plots showing GSEA enrichment score of **(D)** Interferon alpha, **(E)** Interferon gamma, **(F)** TGFβ and **(G)** peroxisome pathway according to MHC status on HRS cells.

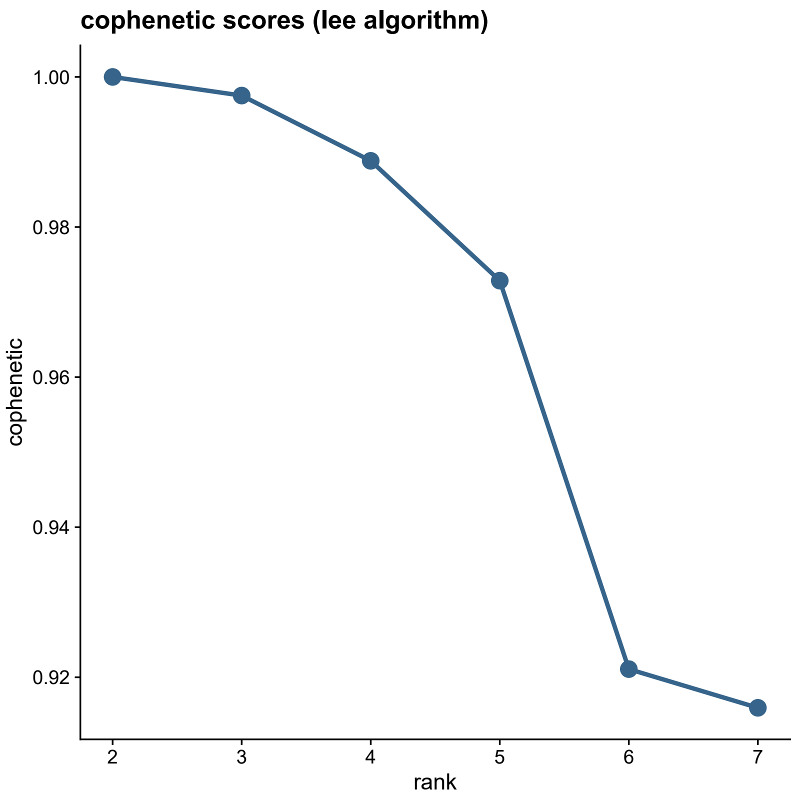


**Figure S10. Cophenetic correlation for increasing number of NMF-defined clusters.**

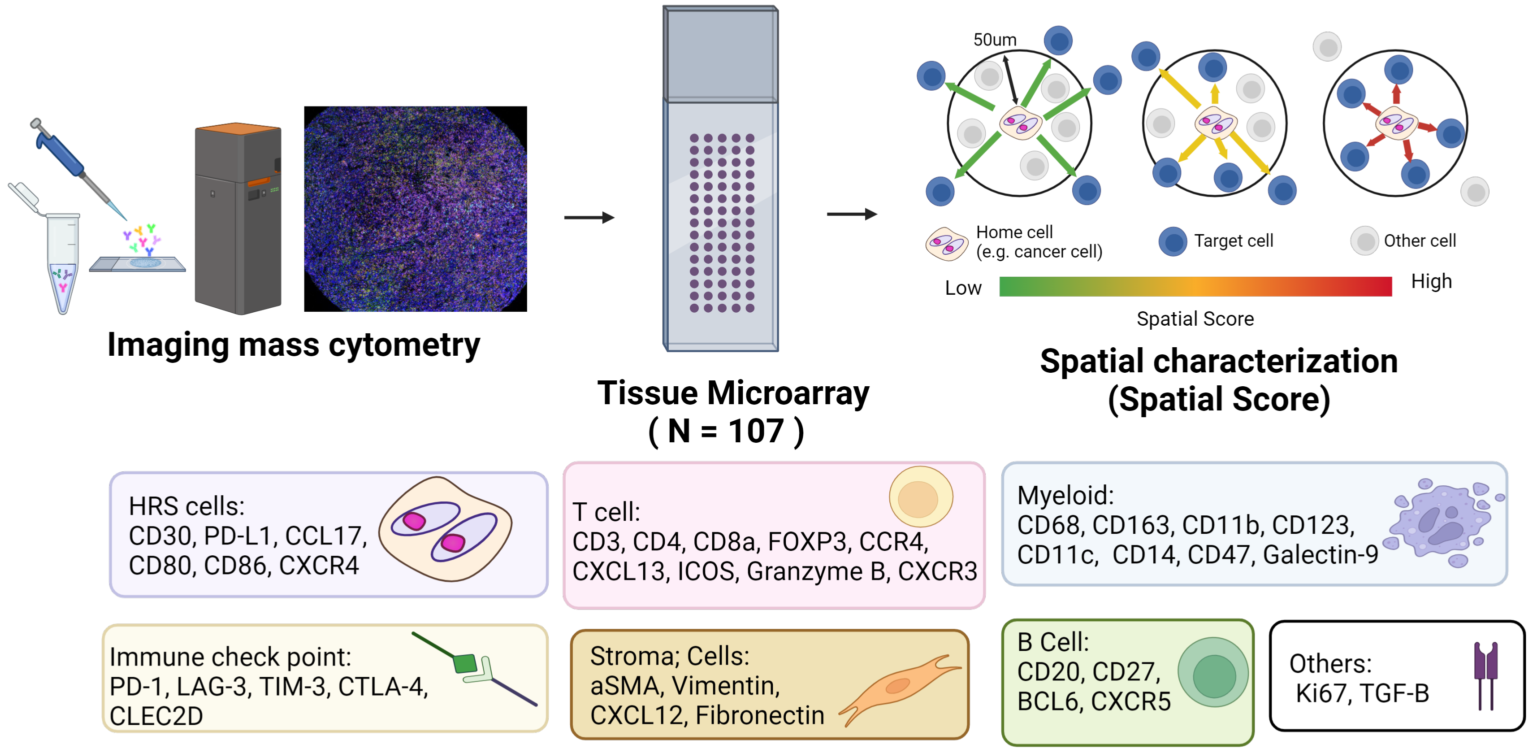


**Figure S11. Overview of the imaging mass cytometry (IMC) analysis and panel.**

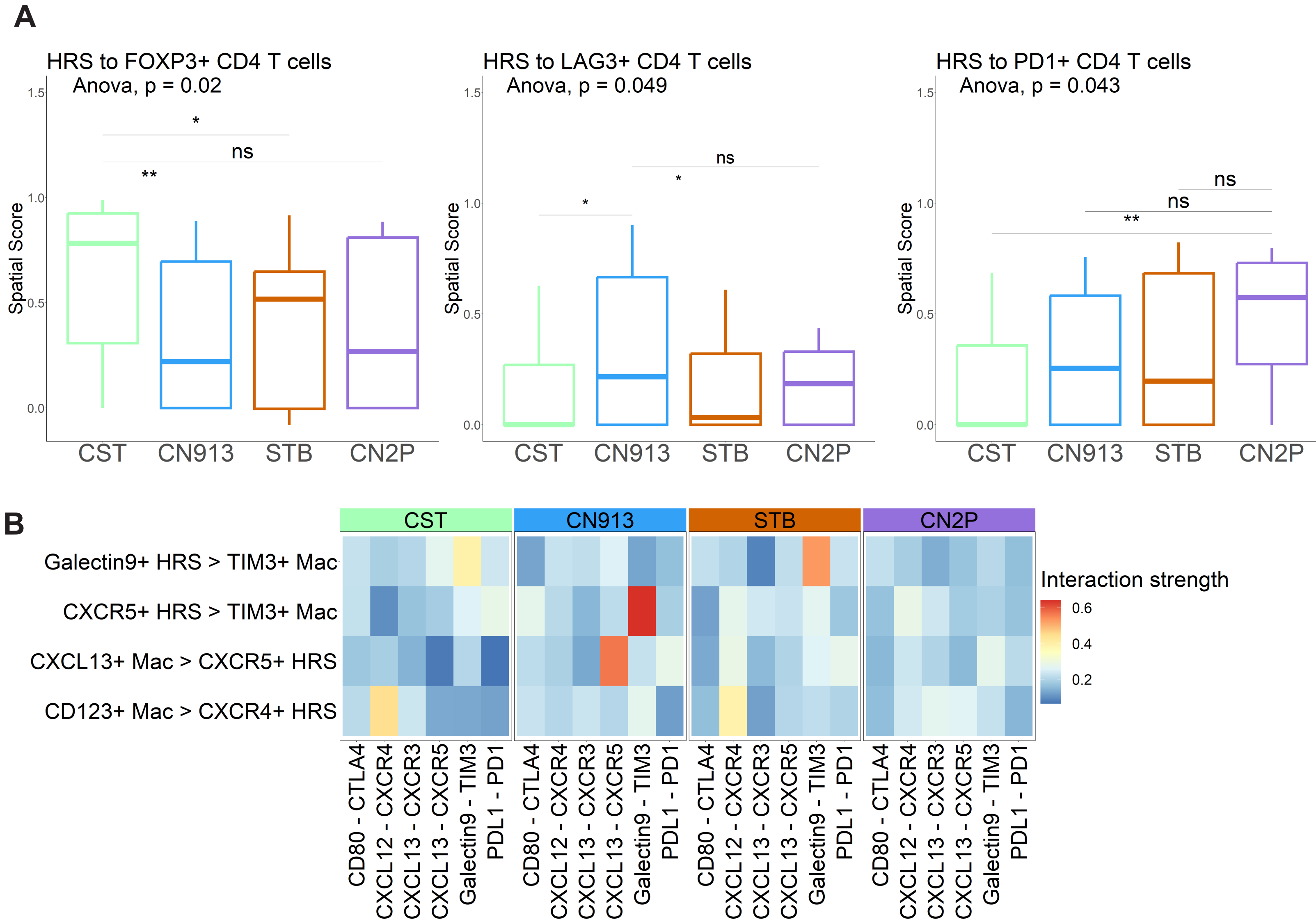


**Figure S12. Spatial tumor-microenvironment correlates with molecular subtype in CHL.** (**A**) Box plot indicating the spatial score for the indicated CD4 T cell subsets, (**Left**) FOXP3+ Treg, (**Right**) LAG3+CD4+ T cells, and (**Bottom**) PD1+CD4+ T cells, near Hodgkin and Reed-Sternberg (HRS) cells according to molecular subtypes. Asterisks represent associated *p*-values (**p*< 0.05; ***p*< 0.01). (**B**) The heatmap shows significant ligand and receptor interaction between HRS cells and macrophage populations using CellChat in each molecular subtype.

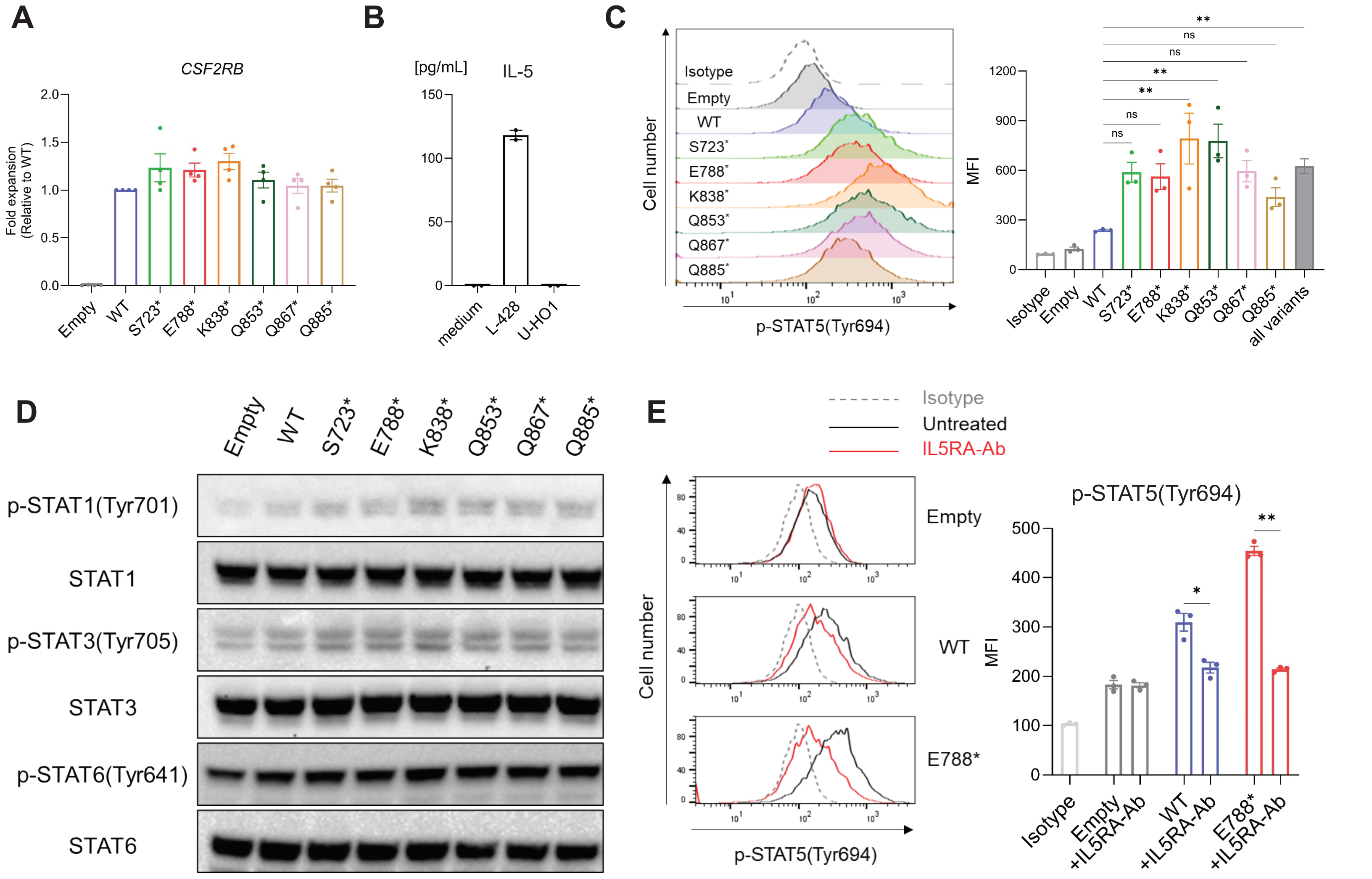


**Figure S13. Functional characterization of *CSF2RB* mutations in the L-428 model.**
**(A)** qPCR analysis of *CSF2RB* expression in the L-428 model. *CSF2RB* expression levels were normalized to *GAPDH* levels, and the figure shows fold expansion in each cell relative to WT cells. Data are presented as mean ± SEM. **(B)** The amount of IL-5 in the supernatant of L-428 and U-HO1 cells by Luminex analysis. Data are presented as mean ± SEM. **(C)** Representative image of p-STAT5 expression by flow cytometry (FCM) analysis (left). Mean fluorescence intensity (MFI) of p-STAT5 expression (right). Data are presented as mean ± SEM. Statistical analyses were conducted using one-way ANOVA followed by Dunnett's multiple comparisons test. **(D)** Western blot (WB) analysis of STAT1, STAT3, and STAT6 expression and their phosphorylation status in the L-428 model. **(E)** p-STAT5 levels by FCM in L-428 cells after the treatment by antibodies targeting surface IL5RA for 3 hours. Representative image of p-STAT5 expression by FCM analysis (left). MFI of p-STAT5 expression (right). Data are presented as mean ± SEM. Statistical analyses were conducted using one-way ANOVA followed by Dunnett's multiple comparisons test.

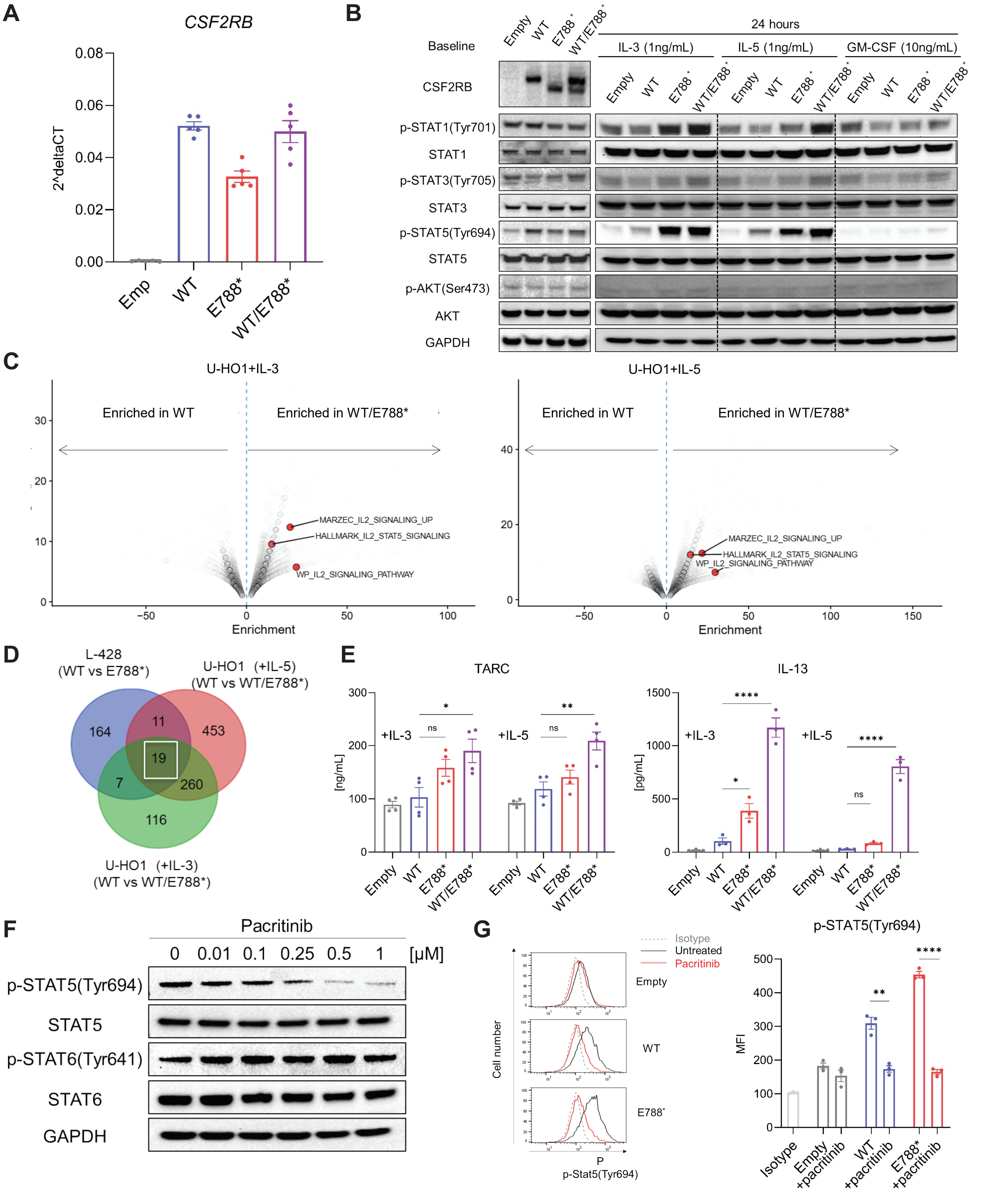

A
CSF2RB
2^deltaCT
0.08
0.06
0.04
0.02
0.00
Emp
WT
E788*
WT/E788*
B
Baseline
Empty
WT
E788*
WT/E788*
24 hours
IL-3 (1ng/mL)
IL-5 (1ng/mL)
GM-CSF (10ng/mL)
CSF2RB
p-STAT1(Tyr701)
STAT1
p-STAT3(Tyr705)
STAT3
p-STAT5(Tyr694)
STAT5
p-AKT(Ser473)
AKT
GAPDH
C
U-HO1+IL-3
U-HO1+IL-5
Enriched in WT
Enriched in WT/E788*
MARZEC_IL2_SIGNALING_UP
HALLMARK_IL2_STAT5_SIGNALING
WP_IL2_SIGNALING_PATHWAY
Enrichment
D
L-428
(WT vs E788*)
U-HO1 (+IL-5)
(WT vs WT/E788*)
U-HO1 (+IL-3)
(WT vs WT/E788*)
164
11
453
19
7
260
116
E
TARC
IL-13
+IL-3
+IL-5
[ng/mL]
[pg/mL]
ns
*
**
****
Empty
WT
E788*
WT/E788*
F
Pacritinib
0
0.01
0.1
0.25
0.5
1
[μM]
p-STAT5(Tyr694)
STAT5
p-STAT6(Tyr641)
STAT6
GAPDH
G
Isotype
Untreated
Pacritinib
Empty
WT
E788*
Cell number
p-Stat5(Tyr694)
p-STAT5(Tyr694)
MFI
Isotype
Empty +pacritinib
WT +pacritinib
E788* +pacritinib

**Figure S14. Functional validation and characterization of intrinsic phenotypes of *CSF2RB* mutations. (A)** qPCR analysis of *CSF2RB* expression in U-HO1 model. *CSF2RB* expression levels were normalized to *GAPDH* levels. Data are presented as mean ± SEM. **(B)**WB analyses of CSF2RB expression and phosphorylation status of STAT, and AKT proteins at baseline, and after cytokine stimulation for the indicated times. **(C)** Pathway enrichment plots between WT versus WT/E788* cells in U-HO1 model after IL-3 (left) or IL-5 (right) stimulation for (box) 24 hours. **(D)** Venn diagram illustrating the overlap of genes with consistent expression changes (absolute fold change ≥ 1.4) across the three experimental models: L-428, U-HO1 + IL-3 (1 ng/mL) for 24 hours, and U-HO1 + IL-5 (1 ng/mL) for 24 hours. **(E)** TARC (left) and IL-13 (right) concentrations in supernatant from U-HO1 model after IL-3 (1 ng/mL) or IL-5 (1 ng/mL) stimulation for 48 hours. Data are presented as mean ± SEM. Statistical analyses were conducted using one-way ANOVA followed by Dunnett's multiple comparisons test per each experiment model. **(F)** WB analysis of phosphorylation status of STAT5/6 after pacritinib treatment for 48 hours in L-428 E788* cells. **(G)** p-STAT5 levels by FCM in L-428 cells after pacritinib treatment for 48 hours. Representative image of p-STAT5 expression by FCM analysis (left). MFI of p-STAT5 expression (right). Data are presented as mean ± SEM. Statistical analyses were conducted using one-way ANOVA followed by Dunnett's multiple comparisons test.

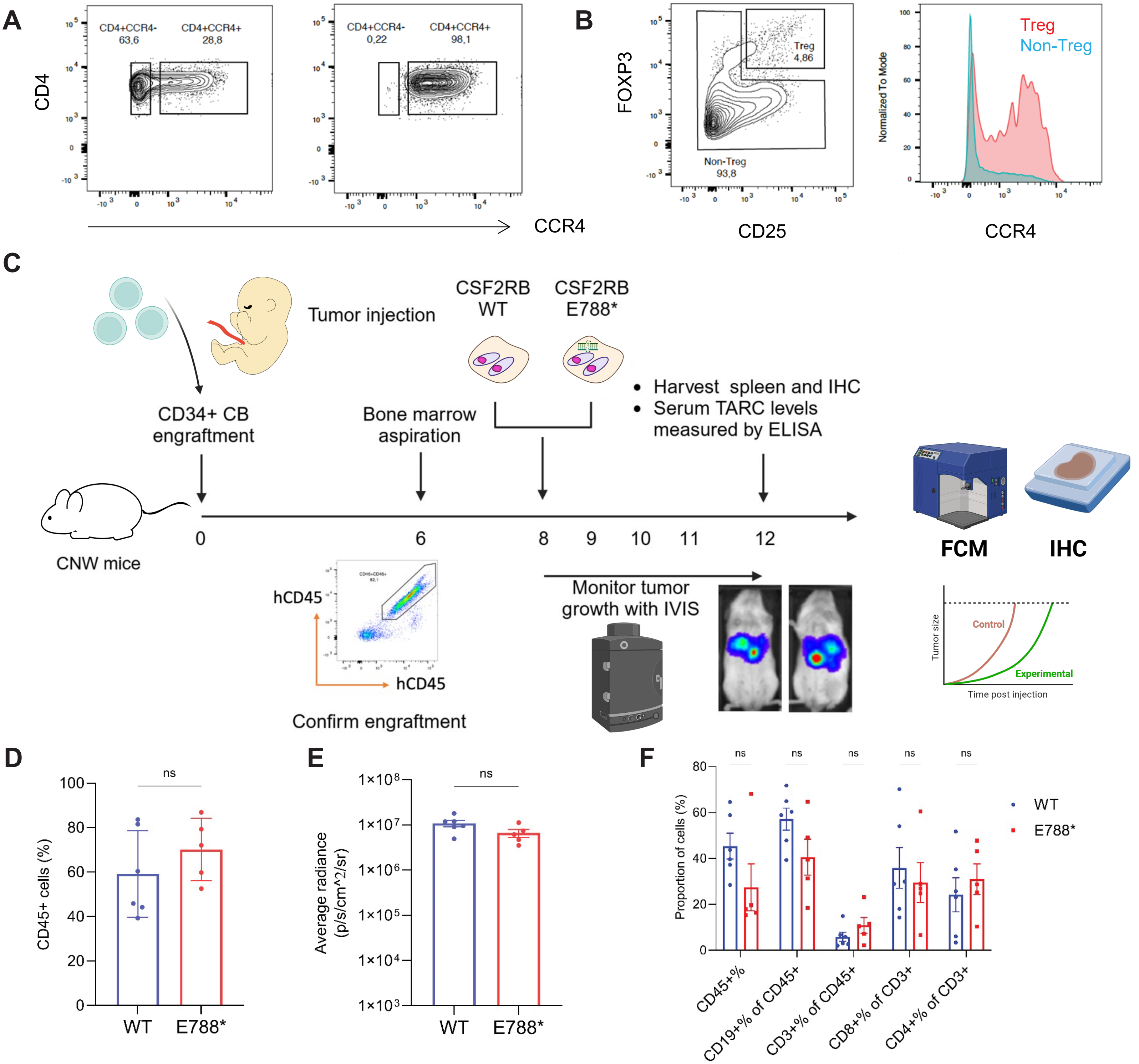


**Figure S15. Characterization of extrinsic phenotypes of CSF2RB mutations. (A)** Representative FCM images showing CD4 and CCR4 expression of CD8-CD3+ healthy donor-derived human PBMCs before (left) and after sorting (right). **(B)** Representative gating strategy for CD4+CD25+FOXP3+ Treg cells and non-Treg cells (**left**), and their CCR4 expression (**right**). **(C)** Experimental schema for humanized xenograft model: CD34+ cord blood (CB) cells were injected intravenously into CNW (NOD.Cg-Rag1nullIl2rgnull/SzJ-W41/41) mice. Engraftment efficiency was measured at week 6 after CB injection. Then, luciferase-labeled L-428 CSF2RB WT or E788* cells were injected intravenously. Tumor burden was monitored by in vivo imaging system (IVIS). **(D)** CD45+ human cells in the bone marrow at week 6 between CSF2RB WT(n=6) and E788* (n=5) groups. An unpaired, two-sided t-test was used to compare the spleen between CSF2RB WT (n=6) and E788* (n=5) groups. The data are shown as mean ± SEM. **(E)** The average radiance in the whole body monitored by IVIS. Unpaired, two-sided t-test was used to compare the spleen between CSF2RB WT (n=6) and E788* (n=5) groups. The data are shown as mean ± SEM. **(F)** Immunophenotyping flow cytometry of spleen identified CD19+ B-cells, CD4+ and CD8+ T-cells. An unpaired, two-sided t-test was used to compare between CSF2RB WT (n=6) and E788* (n=5) groups. Data are presented as mean ± SEM.

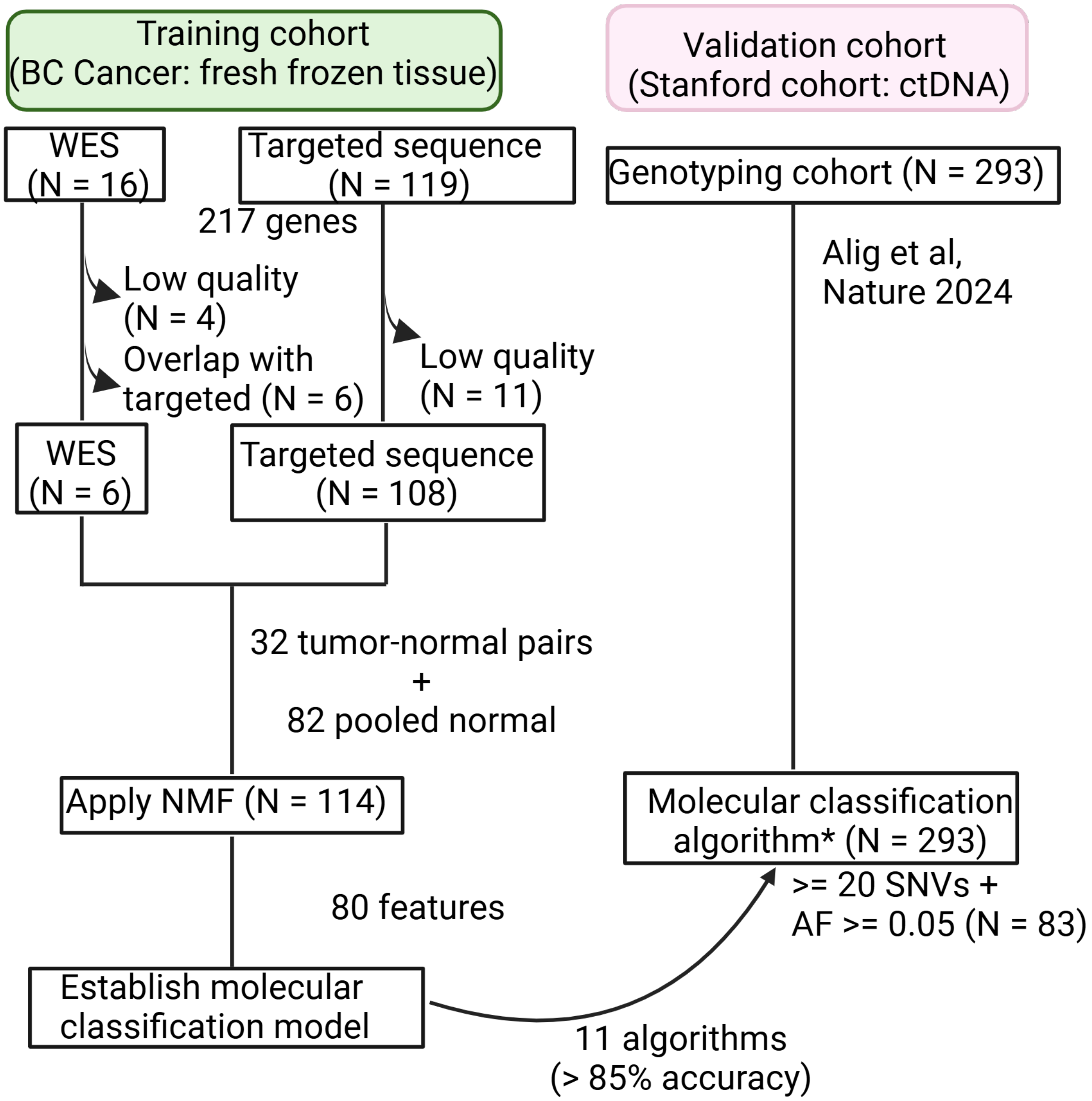


**Figure S16. Cohort and study design overview for the HLGen development.**

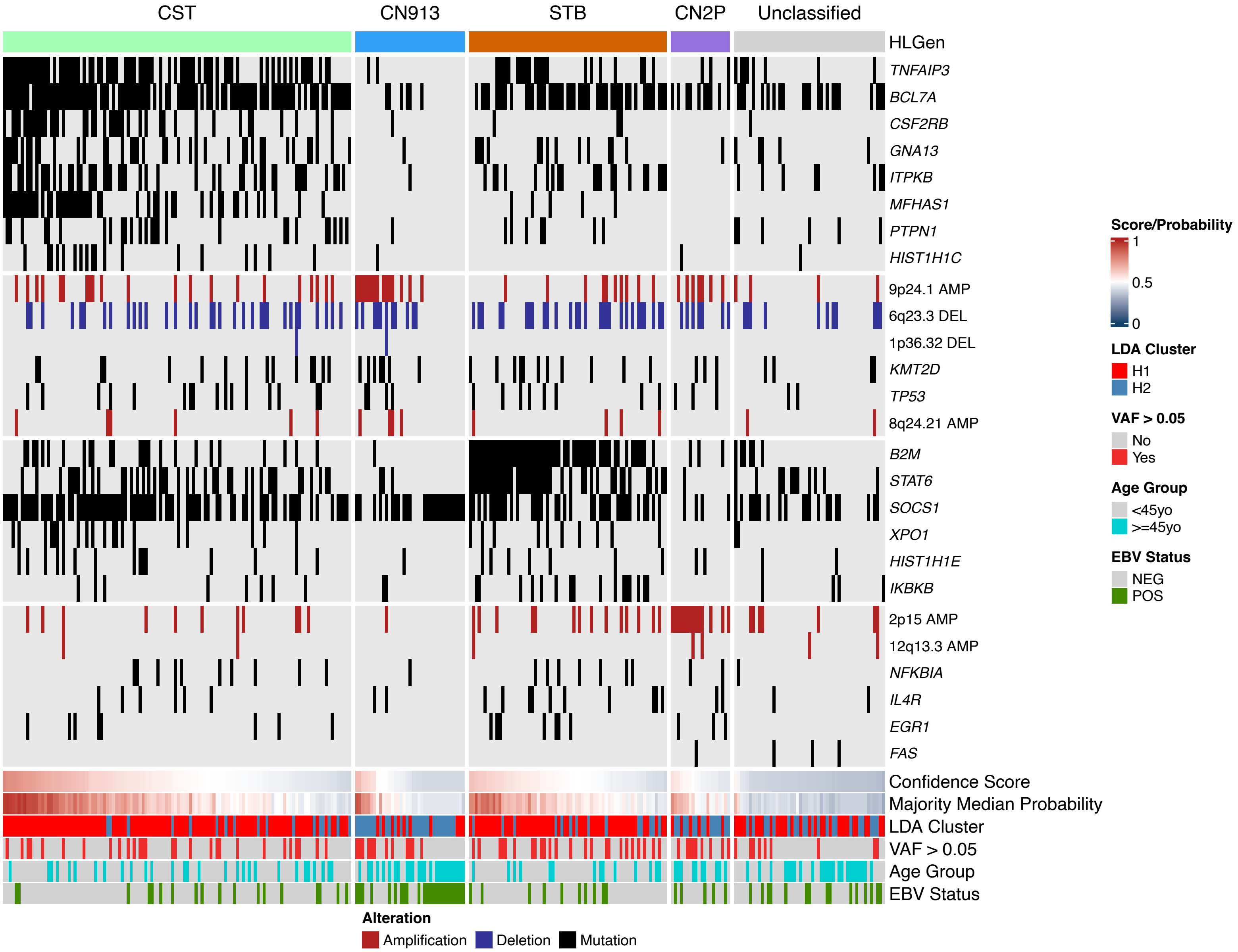


**Figure S17. Oncoplot of the CHL samples from Stanford ctDNA cohort (n = 293), with clinical annotation according to molecular subtype.**

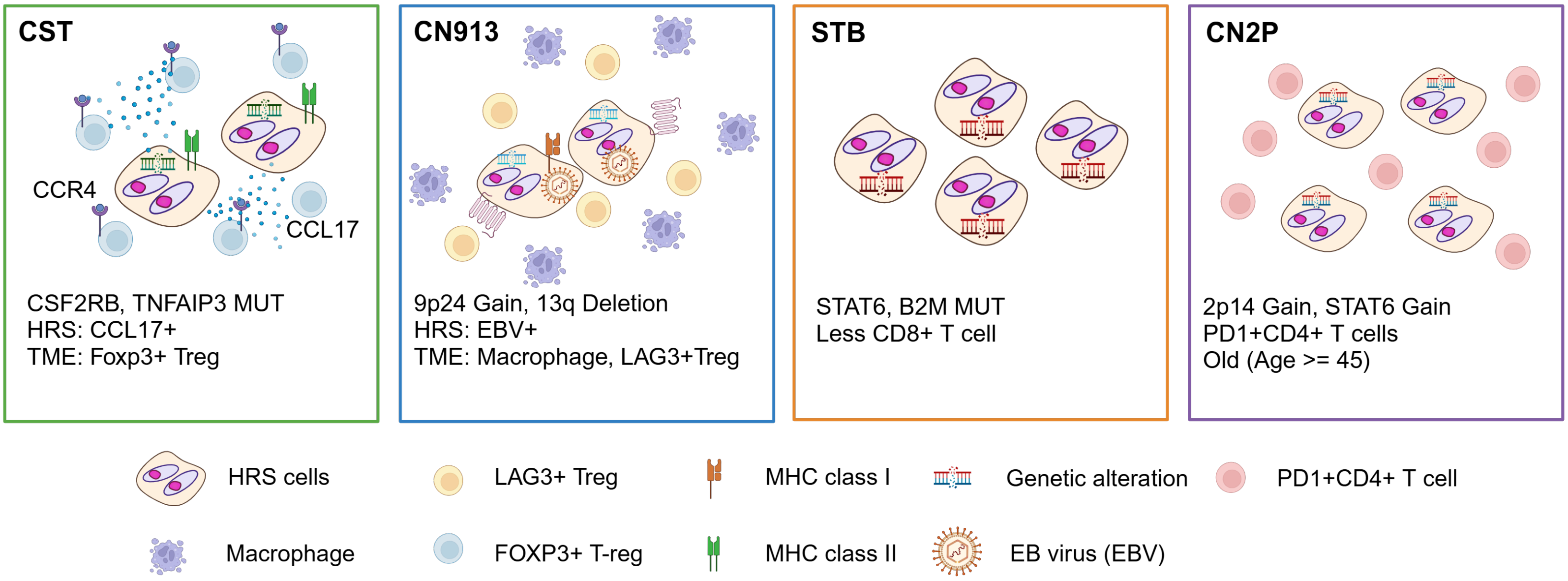


**Figure S18. Graphical summary of molecular subtypes and their clinical and tumor-microenvironment ecosystem correlates in classic Hodgkin lymphoma.**

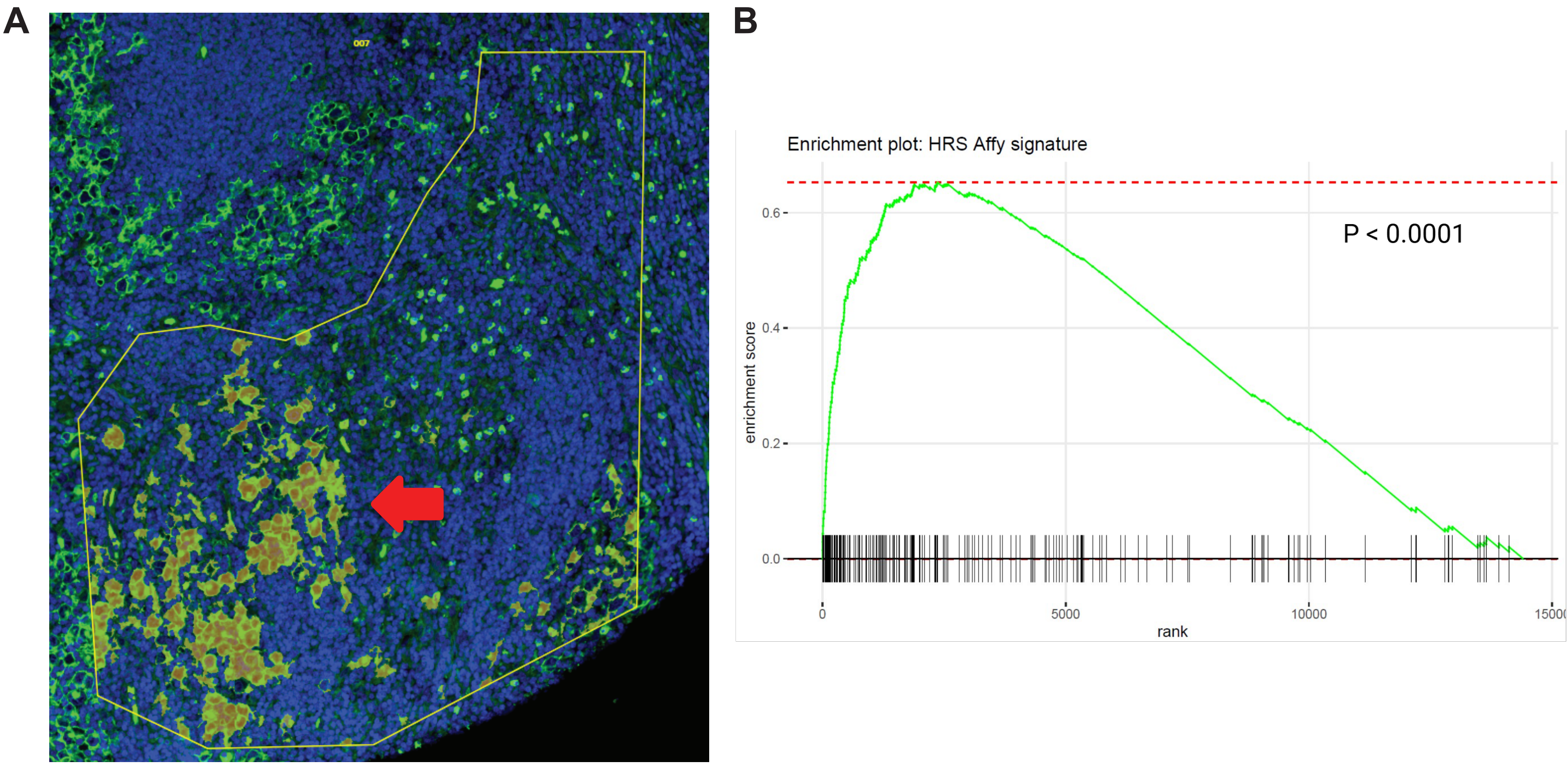


**Figure S19. GeoMx analyses on CD30+ HRS cells. (A)** Zoomed-in ROI showing a representative image of tumor segmented areas (orange), **(B)** In pre-ranked Gene set enrichment analysis (GSEA), gene expression in tumor-segmented area (CD30 high) is significantly and positively correlated with an HRS signature (*p*<0.0001).defined by Affymetrix gene expression data generated from laser microdissected HRS cells (Steidl et al, Blood 2012).

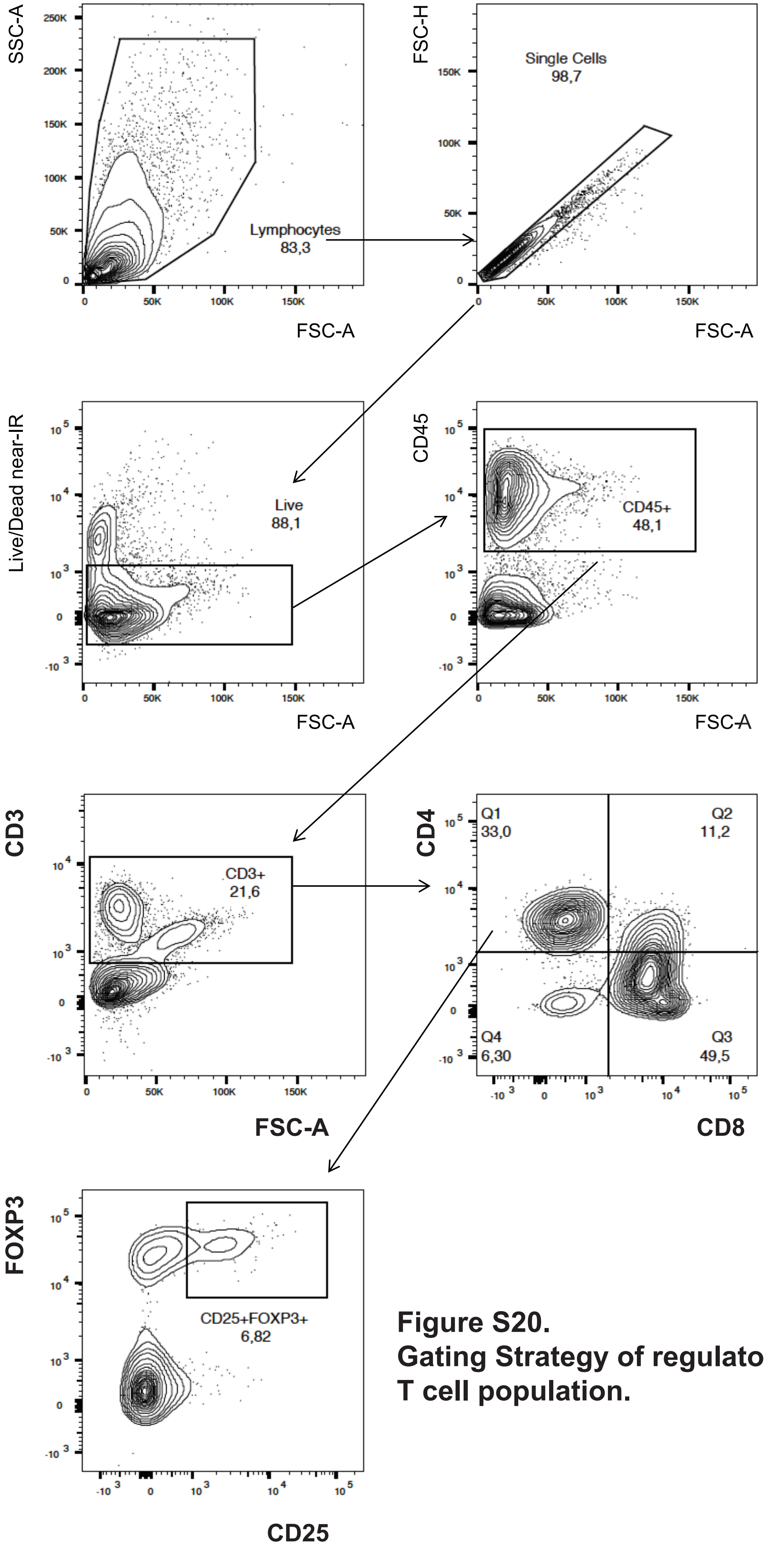


**Figure S20.**
**Gating Strategy of regulatory T cell population.**
